# *Plasmodium falciparum* DNA repair dynamics reveal unique roles for TLS polymerases and PfRad51 in genome diversification

**DOI:** 10.1101/2025.04.07.647301

**Authors:** Akshay Vishwanatha, Xu Zhang, Yi Jing Liu, Annie Leung, Mikayla Herring, Amanda Chan, Susanah Calhoun, Kirk Deitsch, Laura Kirkman

## Abstract

The haploid malaria parasite, *Plasmodium falciparum,* evolved a unique cohort of DNA repair pathways enabling the parasite to survive in a vertebrate host red blood cell and the mosquito vector. *P. falciparum* chromosomes are partitioned into a highly conserved core genome and remarkably diverse, largely subtelomeric regions that contain genes encoding important parasite virulence factors. The molecular mechanisms that maintain this chromosomal structure have not been identified. Here, we describe specific DNA repair pathways that differentiate between hypervariable subtelomeric and conserved core regions of the genome. Based on our previous work, we hypothesized that there are potentially important interactions between translesion (TLS) and homologous recombination (HR) pathways for the diversification of multicopy gene families in *P. falciparum.* Thus, we created knockout parasite lines of the DNA repair enzymes: *PfRad51* and the TLS polymerases *PfPol*ζ and *PfRev1*. We identified that irradiation hypersensitivity varied across the cell cycle for TLSΔ parasites and was uniform across the erythrocytic cycle for *PfRad51*Δ parasites, highlighting the variable roles of these pathways. However, important interactions between these pathways were found when we studied directed double strand break (DSB) repair, which revealed a difference in the DNA damage response according to chromosomal location. *PfRad51* was essential for HR-mediated repair in the core genome. In contrast, we identified a Rad51 independent homology-directed repair in all three of our knockout lines when a DSB was made in the subtelomeric region of the chromosome. We propose that this differential DNA damage response maintains the distinction in diversification across the chromosome.

## Introduction

Malaria inflicts a tremendous health and economic burden, with an estimated 263 million malaria cases in 2023 (an increase of 11 million compared to 2022) and a death toll of approximately 0.5 million globally.^1^ The emergence of resistance to artemisinin and partner drugs used in combination therapy to treat malaria poses a significant risk to global efforts to control the disease ^1^. Malaria is caused by infection with the unicellular obligate eukaryotic parasite of the genus *Plasmodium*. *Plasmodium falciparum* is the most virulent human infectious species. Unique features of the parasite’s genome have been associated with virulence and its propensity to develop drug resistance. *P. falciparum* has a distinctly skewed genome with 80 % AT content, notable repetitive low complexity regions, and a propensity to accumulate small indels during mitotic division^2^. The parasite has a haploid genome for most of its lifecycle, including all stages in the human host, with a brief diploid stage in the mosquito vector. *P. falciparum* has 14 chromosomes displaying a high degree of genome plasticity, mostly confined to the subtelomeric regions that contain multicopy gene families that encode important virulence factors and variant surface antigens. These subtelomeric regions can extend to ∼100 kb in length from the chromosome ends and include telomere-associated repeat elements (TAREs) and members of several hypervariable gene families such as *var*, *rifins* (repetitive interspersed family), *stevor* (subtelomeric variant open reading frame), and *Pfmc-2TM* (*P. falciparum* Maurer’s cleft - 2 transmembrane domain protein)^3,4^. Each family contains multiple genes that display a high degree of sequence variation within the genome of a single parasite and between different geographical isolates^4–6^. We propose that in malaria parasites, DNA repair mechanisms play an important role in genetic diversification, particularly the diversification of the antigenic variant, erythrocytic membrane protein 1 (PfEMP1), encoded by the multicopy gene family *var,* which is present as clusters in the subtelomeric and several internal chromosomal regions (Supplementary Fig 1)^3^. Previous reports highlighted the high variability of these regions compared to the core genome^4^ and possible mechanisms such as non-allelic homologous recombination (NAHR), alternate end joining (Alt-EJ), gene conversion, and telomere healing (TH) contribute to this diversity during the asexual erythrocytic stages^7–10^. Therefore, understanding these different DNA repair pathways in the parasite will provide insight into the complex process of genome diversification, including this critical gene family of virulence factors.

*P. falciparum* possesses a complex yet unique genome maintenance system crucial for survival within the host. DNA repair mechanisms are required to overcome DNA damage induced by genotoxic stress, external damaging agents like mutagens, ionizing radiation, and drugs, including various antimalarials^11–14^. As the parasite proceeds through the intraerythrocytic development cycle (IDC), it undergoes schizogony, where one parasite replicates asynchronously to form 30-50 daughter cells. This development requires multiple rounds of DNA replication; thus, the parasite undergoes significant replication stress. Damaged bases or mistakes during DNA replication result in replication fork stalling. Hence, DNA damage must be repaired for replication to proceed. In addition to various repair mechanisms to remove damaged or mismatched bases, cells have developed specific processes that tolerate mistakes and allow replication to proceed. These processes are called replicative DNA damage bypass pathways or translesion synthesis (TLS)^15^. TLS is a highly conserved process mediated by specialized polymerases that replicate opposite and past damaged bases with lower fidelity, called TLS polymerases. The lower fidelity of these polymerases carries an increased risk of mutagenesis, which can be detrimental but also serve as a source of genomic diversification^15,16^. We recently reported the potential importance of TLS components in diversifying multigene families through ectopic recombination^17^. Rodent malaria parasites also harbor diverse multicopy gene families; however, there is a high degree of identity when comparing sequences between different isolates^18^. This high degree of sequence identity contrasts with much higher diversity between *P. falciparum* isolates. Of note, rodent parasites also lack the machinery for TLS^17^. In the human malaria parasite, components of the TLS complex identified in silico include the scaffold protein subunit *PfRev1* and the catalytic subunit *PfPol*ζ/*Rev3*, with two potential HORMA domain containing Rev 7 orthologs (Fig 1A).

**Figure 1:**
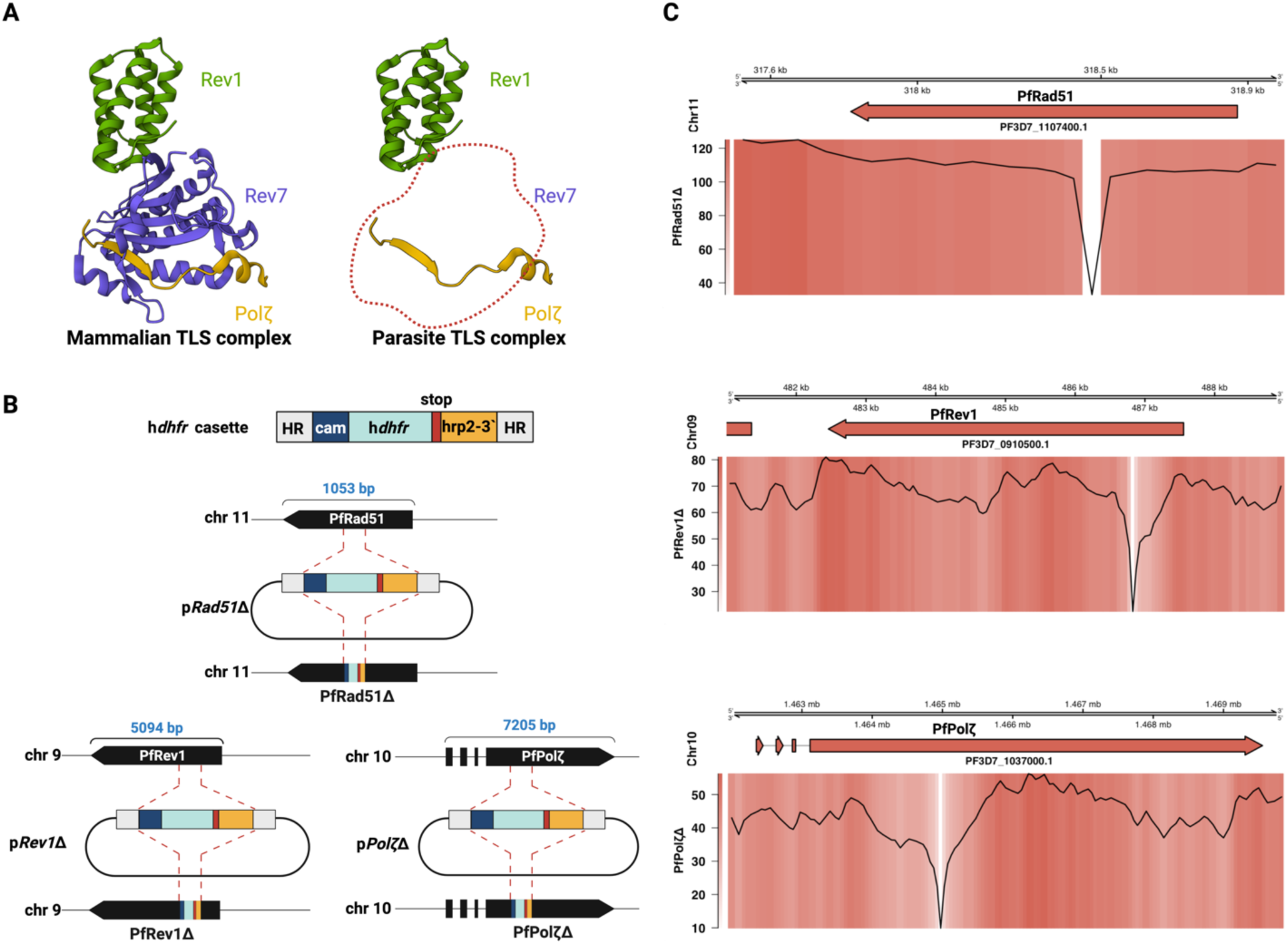
Construction and confirmation of knockouts of *PfRad51*, *PfRev1,* and *PfPol*ζ. (A) Comparison between mammalian and parasite TLS complex (PDB:3VU7). The red dotted line indicates that the Rev7 subunit has not been identified to date in parasites. (B) Overview of the molecular approach used for creating the knockouts of *PfRad51*, *PfRev1,* and *PfPol*ζ. The top cartoon shows the hDHFR (Dihydrofolate Reductase) cassette used for integration and selection. (C) Coverage plots of Nanopore sequencing results of the *PfRad51*Δ, *PfRev1*Δ, and *PfPol*ζΔ. The top panel denotes the genome region, chromosome, and gene. The bottom panel has both heatmap and lineplot to show the integration site where the hDHFR cassette has been integrated, which can be seen as a drastic drop in the coverage (Large dip in the lineplot coinciding with the white region in the coverage plot).

Another lethal type of DNA damage is double-strand breaks (DSBs), which are repaired mainly by the non-homologous end joining (NHEJ) and the homologous recombination (HR) repair pathways. *P. falciparum* and all closely related blood-borne apicomplexan parasites lack the components of canonical NHEJ, making HR the major pathway for repairing DSBs^17,19^. Since Plasmodium is haploid throughout its IDC and lacks homologous pairs of chromosomes, it is surprising that it depends on HR for DSB repair. A relatively inefficient alt-EJ pathway was identified that might constitute a method of repairing DSBs in the absence of homologous templates^20,21^. In addition, we and others have reported how antigen diversification occurs through mitotic recombination mediated by HR in *P. falciparum* ^7–10^. A defining feature of HR is the utilization of homologous DNA sequences as a repair template where the broken DNA molecule finds and invades the homologous DNA molecule. The Rad51 recombinase, assembled on the ssDNA at the site of the break as an oligomeric nucleoprotein filament, performs this function^22^. Therefore, in *P. falciparum*, HR is the main pathway by which DSBs are repaired, followed by alt-EJ in the absence of homologous templates.

Based on data suggesting the importance of the TLS polymerases and the HR pathway in the maintenance and diversification of the parasite genome, we set out to determine the general function of these proteins as well as their specific role in genome diversification.

## Results

### Creation of homologous recombination and translesion synthesis knockout mutants

To study the function of *PfRad51* (PF3D7_1107400) and TLS polymerases *PfRev1* (PF3D7_0910500) and *PfPol*ζ (PF3D7_1037000) in *P. falciparum*, we first tested if these genes were essential for parasite viability. Rad51 is especially interesting because it is essential for survival in vertebrates where null mutation of Rad51 leads to embryonic lethality and death of mammalian cell lines^23^, whereas Rad51 is not essential in other systems such as budding yeast^24^, *T. brucei* ^25^, fission yeast *S. pombe*^26^ and *S. japonicus* ^27^. First, we generated a functional knockout of *PfRad51* using CRISPR/Cas9 (Clustered Regularly Interspaced Short Palindromic Repeats/CRISPR-associated protein-9 nuclease). As shown in Fig 1B (top cartoon), a repair block containing the selection marker hDHFR, encoding resistance to the drug WR99210 (WR) with flanking homologous regions upstream and downstream of the selected PAM (protospacer adjacent motif) site in the target gene, *PfRad51*, constituted the donor plasmid^28^. The donor plasmid also contained the single guide targeting the sequence coding for the ATPase domain of the protein (TTATTTGGTGAATTTCGTAC|AGG). A similar strategy was used to generate knockouts of the TLS polymerases *PfRev1* (ATATTAATAGTTCAAAACGG|AGG, BRCT domain) and *PfPol*ζ (TTATCTTTTTAATGAAAGTG|AGG, upstream of DNA-directed DNA polB domain) using CRISPR/Cas9 as outlined in Fig 1B. Rev1 is not essential for the survival of higher eukaryotes, while Polζ is essential during development in mammalian systems^22,29,30^ whereas both genes are non-essential in yeast^31^. The generated donor plasmid (Supplementary Table 1) along with a Cas9 plasmid (pUF1-Cas9)^28^ were co-transfected into a wildtype (WT) 3D7 background to generate the three knockout mutants - *PfRad51*Δ, *PfRev1*Δ, and *PfPol*ζΔ. The successful transfections and comparable growth rates of the mutant lines to WT 3D7 line indicate that in *P. falciparum,* these three genes are not essential for survival (Supplementary Fig 2). The knockouts were confirmed by Illumina whole genome sequencing, as shown in Fig 1C. The loss of the part of the gene replaced by the selection cassette is represented by the sudden loss of coverage towards the N-terminal region. Our attempts to generate double knockouts of *PfRad51* in each of the TLS polymerase knockout backgrounds did not yield viable parasites, indicating the essentiality of TLS polymerases in the absence of HR or that *PfRad51*, *PfRev1*, and *PfPol*ζ function in parallel repair pathways.

### *P. falciparum* TLS polymerases are essential for the survival of irradiated ring-stage parasites

The IDC of *P. falciparum* begins with the invasion of a red blood cell (RBC) by a single merozoite containing one copy of each chromosome. This merozoite develops into the ‘ring’ form from 0 h to 24 h post-invasion. At this stage of the IDC, the parasite contains a single copy of the genome. DNA replication at later stages of the cycle results in multiple copies inside an intact nucleus, potentially providing a template for the HR pathway to repair DSBs (Fig 2A)^32–35^. Therefore, in ring-stage parasites, given the absence of NHEJ in Plasmodium, there is greater reliance on alternate DSB repair pathways given a lack of template for repair required by HR. Different pathways are likely responsible for DNA repair in the ring and late IDC stages. Thus, sensitivity to DNA damage would also vary depending on the parasite’s position within the replicative cycle^3,35^. Therefore, the knockouts of *PfRad51* and the TLS components described in the previous section were tested for their sensitivity to DNA damage using X-rays as a source of ionizing radiation (Fig 2B). Ionizing radiation such as X-rays is known to induce multiple kinds of DNA damage including base damage, single-strand breaks (SSB), double-strand breaks (DSB) that require different repair pathways such as nucleotide excision repair (NER), base excision repair (BER), mismatch repair (MMR) and DSB repair pathways for the exposed cell to recover ^36^.

**Figure 2:**
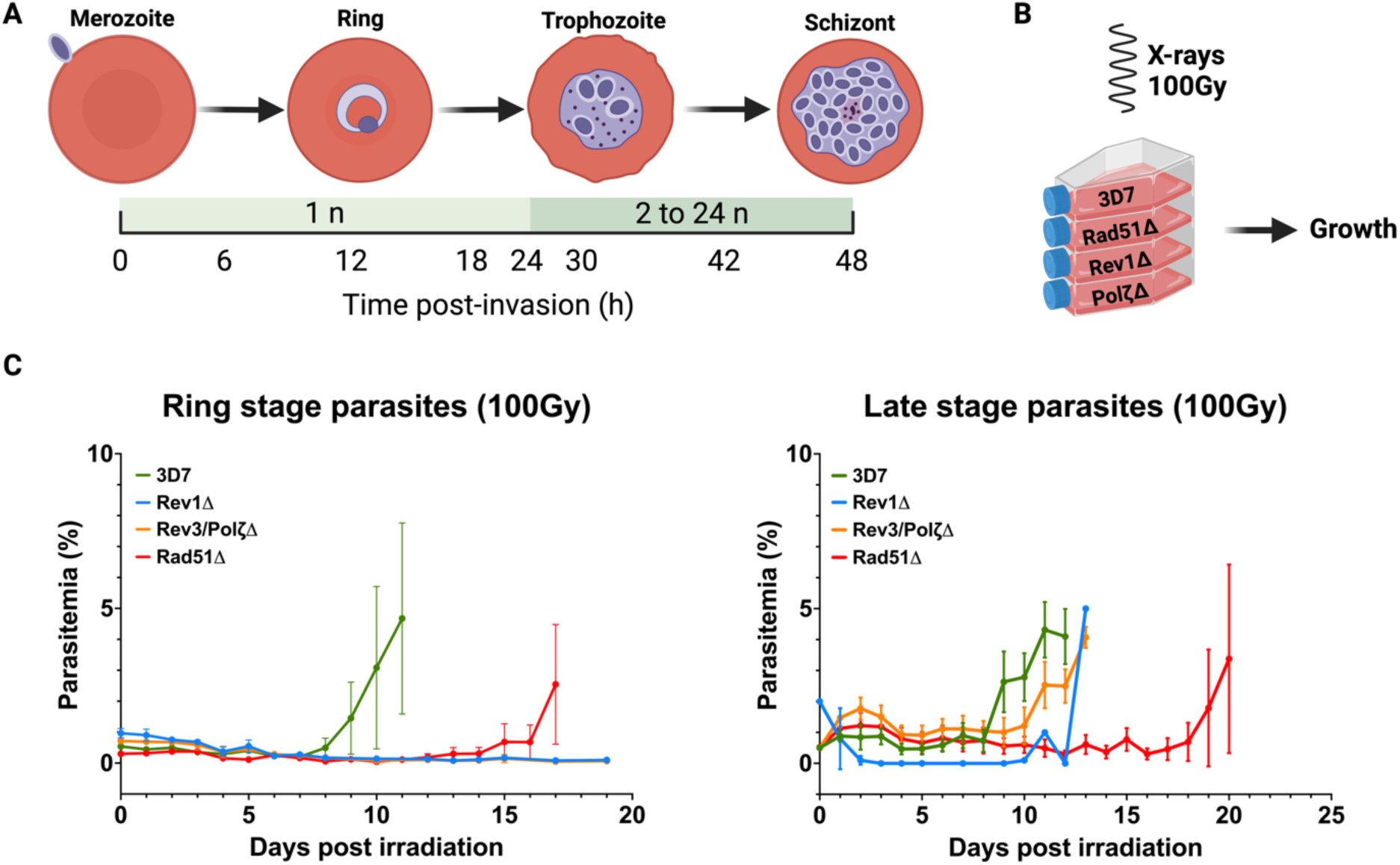
Growth analysis of knockout cell lines. (A) The top panel represents the main stages of the intraerythrocytic developmental cycle (IDC) of *P. falciparum*. The bottom panel shown the change in DNA content over the different stages of the IDC tracked as hours post-invasion (hpi). (B) Overview of the irradiation exposure of the 4 different cell lines - WT, *PfRad51*Δ, *PfRev1*Δ, and *PfPol*ζΔ. (C) Growth curve of 100 Gy irradiated ring stage (0-24 hpi) and late stage (24-48 hpi) parasites.

*PfRad51*Δ, *PfRev1*Δ, and *PfPol*ζΔ, as well as WT 3D7 parasites were exposed to 100 Gy irradiation at both the ring- and late-stage of the replicative cycle. Cultures were then monitored for up to three weeks to determine how the loss of different DNA repair proteins affected the parasites’ ability to repair DNA damage and reestablish growth. For both ring- and late-stages, WT 3D7 parasites recovered from an initial growth inhibition within 9 days of irradiation exposure (Fig 2C, green line), thereby establishing the time required for recovery when all repair pathways are active. Given that malaria parasites are thought to depend almost exclusively on HR for repairing DSBs, and since Rad51 is required for HR in model organisms, we anticipated a near-complete elimination of recovery in the *PfRad51*Δ line. We also predicted that the loss of Rad51 might cause a more pronounced effect in late-stage parasites, which contain multiple copies of the genome, and thus, DSB repair would be further dominated by HR. Remarkably, both ring- and late-stage parasites were able to recover from irradiation in the absence of Rad51 (15 days for ring-stage parasites vs. 19 days for late stages), and the loss of *PfRad51* did not appear to be associated with a substantial difference in survival between these two stages (Fig 2C, red line). These data indicate that despite the presumed dependence on HR for DSB repair in malaria parasites, sufficient alternative repair activity exists to enable reproducible recovery from extensive DNA damage without Rad51. Our previous work indicated that at this irradiation dose, virtually all parasites sustain damage and those parasites that survive display longer telomeres, a hallmark of recovery from DNA damage^37^.

We then examined the effect of the loss of TLS polymerases on parasite recovery from the same irradiation dose. We observed that both *PfRev1* and *PfPol*ζ are essential for the survival of ring-stage parasites when irradiated (Fig 2C, blue and orange lines) as the knockouts fail to recover after more than 20 days. The essentiality of these two TLS polymerases, even when the canonical HR pathway is intact, suggests they function in a Rad51-independent repair pathway required for DNA repair in haploid, ring-stage parasites. Interestingly, the loss of TLS polymerases had little effect on the parasites’ ability to recover from irradiation in later stages, when PfRad51-mediated HR presumably dominates (Fig 2C, blue and orange lines). These observations demonstrate distinct DNA damage sensitivity for our different knockout parasite lines.

### *PfRad51* and PfTLS polymerases differentially impact SNP and indel accumulation

To further validate the canonical functions of TLS polymerases and Rad51, we examined the genome sequence of parasites that had recovered from irradiation (Fig 3A). Briefly, we irradiated WT 3D7, *PfRad51*Δ, *PfRev1*Δ, and *PfPol*ζΔ late-stage parasites with 100 grays of X-rays three times, each time letting the parasites recover. Clonal lines were established from the recovered parasites, and genomic DNA was isolated, purified, and Illumina sequenced (NovaSeq 6000, Kapa HiFi library kit); the sequencing data were processed to identify single nucleotide polymorphisms (SNPs), insertions and deletions (Indels) using Freebayes ^38^ and SnpEff ^39^ as outlined in Fig 4A.

**Figure 3:**
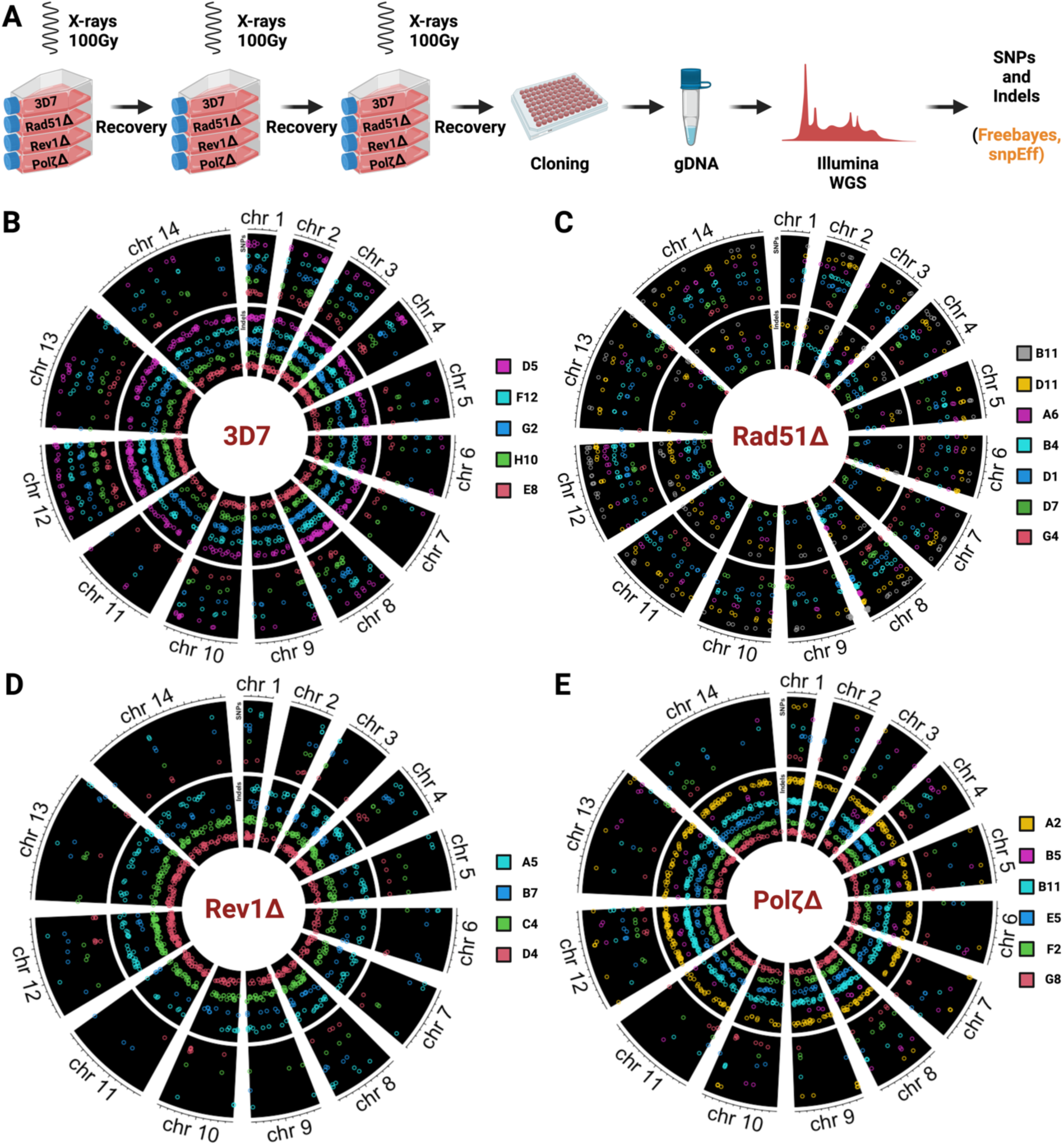
Irradiation induced SNPs and Indels. (A) Overview of irradiation experiment to analyze SNPs and Indels using Illumina whole genome sequencing. (B-E) Circos plots of SNPs (outer arcs) and Indels (inner arcs) distributed throughout the genome of *P. falciparum* cell lines – (B) WT-3D7, (C) *PfRad51*Δ, (D) *PfRev1*Δ, and (E) *PfPol*ζΔ after 3x irradiation with 100 Gy. The different colors represent the clones, and the arcs represent the chromosomes.

**Figure 4:**
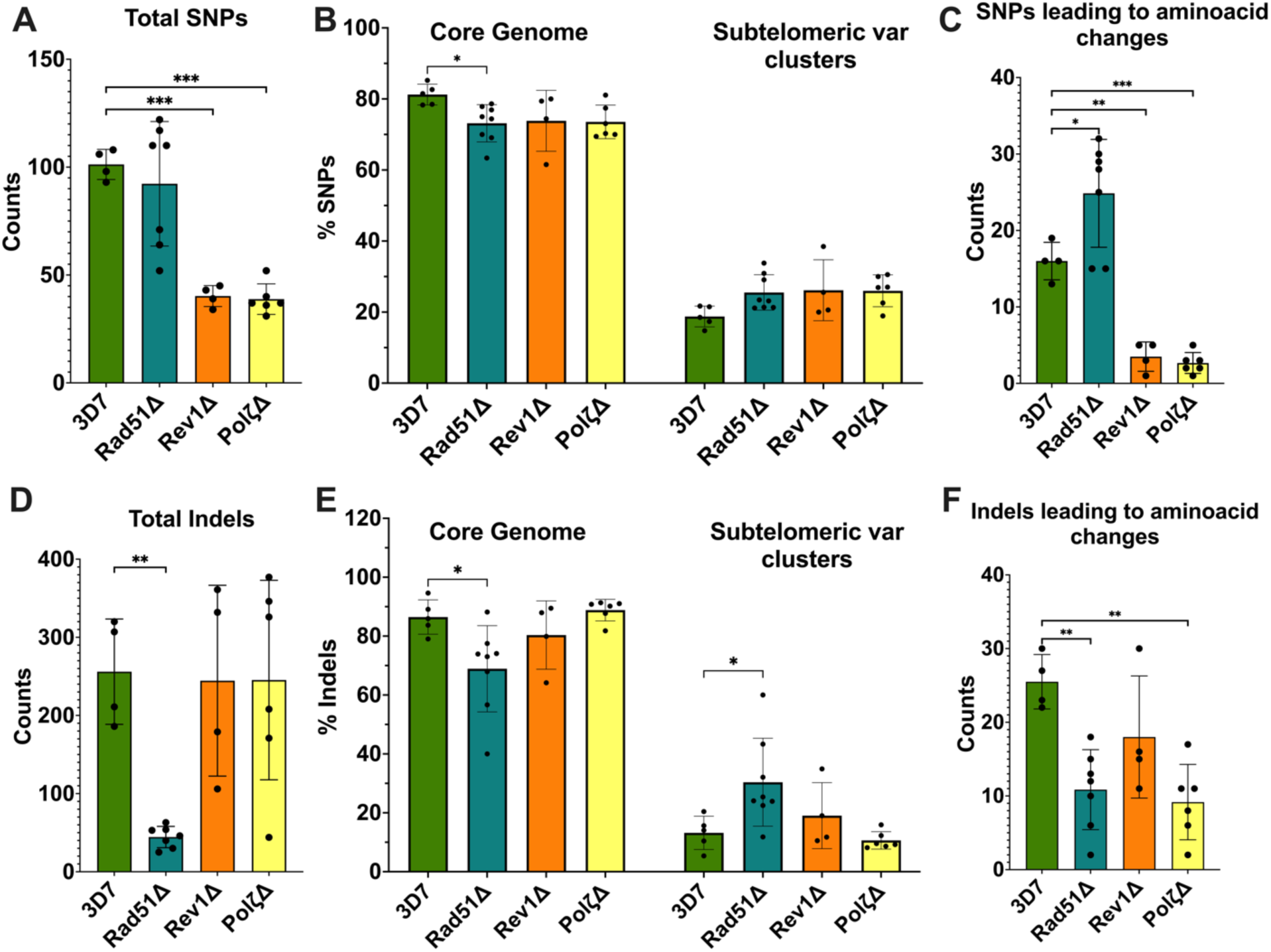
Analysis of SNPs and Indels determined from. Figure 3: Quantification of total SNPs (A) and Non-synonymous SNPs (C); Quantification of total Indels (D) and Non-synonymous Indels (F) from Figure 3. Statistical significance was determined using ordinary one-way ANOVA followed by Šídák’s multiple comparisons test *(*p*=0.05-0.01), **(*p*=0.01-0.001),***(*p*=0.001-0.0001). Distribution of SNPs (B) and Indels (E) across the core genome and the *var* clusters. Statistical significance was determined using Two-way ANOVA followed by Dunnett’s multiple comparison test *(*p*=0.05-0.01), **(*p*=0.01-0.001), ***(*p*=0.001-0.0001).

To determine the effect of the loss of TLS polymerases and Rad51 on the accumulation of mutations in *P. falciparum*, we analyzed the complete genome sequences of irradiated parasites from multiple clones from each knockout line and compared them to WT 3D7 parasites. As expected, the PfTLS knockout lines display a hypomutable phenotype (Fig 3D and E, 4A), with the WT 3D7 more prone to accumulating SNPs post irradiation (Fig 3B). The distribution of repaired damage was the same across the chromosome for all lines tested, with most SNPs and indels occurring in the core genome when looking at the percent of mutations (Fig 4B, E). The PfTLS knockout mutants accumulated considerably lower non-synonymous SNPs, indicating that any damage in coding regions was not repaired and, thus likely lethal (Fig 4C). In the case of indels, which are thought to originate from DSBs^40^, disruption of the PfTLS pathway had little effect (Fig 4D). Although there are reports of the involvement of the TLS pathway in DSB repair in other systems^41–43^, *PfRev1*Δ and *PfPol*ζΔ show identical phenotypes where the total number of indels is unchanged compared to WT 3D7 (Fig 3D and E, 4D). Interestingly, both *PfRev1*Δ and *PfPolζ* Δ display a trend toward decreased accumulation of non-synonymous indels, which reaches statistical significance in the case of *PfPolζ*Δ (Fig. 4F), suggesting a greater role for TLS polymerases in repairing DNA damage within coding regions. Coding regions, specifically transcribed regions of the genome, can be more vulnerable to DNA damage due to the topological strain from transcription and a more open chromatin structure, perhaps leading to increased accumulation of DSBs. In *P. falciparum*, coding regions also have a pronounced increase in GC content compared to intergenic regions, which could affect DNA repair. Polζ can play a role in repairing DSBs through a post-replicative DNA recombination pathway observed in other model systems^44^. In addition, Polζ is enlisted as a DNA polymerase during the recombination process, and the action of Polζ leads to mutations in yeast ^41^. Therefore, it is possible that in *P. falciparum*, the TLS polymerases might also have important cross functionality responding to the repair of excised damaged bases as well as DSB repair from stalled replication forks or direct DNA damage.

The loss of *PfRad51* did not affect the frequency and number of SNPs across the whole genome (Fig 3B, Fig 4A). We did, however, observe a small but statistically significant increased incidence of non-synonymous SNPs (Fig 4C, Supplementary Fig 3A), suggesting that in the absence of Rad51, repair in the coding regions is error-prone. In the case of indels, both the total number of indels and non-synonymous indels were drastically reduced (Fig 4D and F). This result indicates that Rad51 plays an important role in the repair of damage across coding and noncoding regions with different outcomes (an increase in SNP accumulations and a decrease in indels), supporting that different compensatory pathways respond to different types of damage in the absence of Rad51. This skew towards the accumulation of mutations in parasite coding regions was also seen in other work where DNA replication or repair proteins were modified or deleted^45,46^.

### *P. falciparum* exhibits differential DNA repair in the core genome and subtelomeric regions

The Plasmodium core genome and the subtelomeric regions are considerably different in terms of sequence diversity. While the core genome is largely conserved when comparing different isolates, the subtelomeric regions are highly diverse, suggesting a greater potential for recombination within these regions^4^. Previous work analyzing parasites grown under standard culture conditions found that most small indels occurred in noncoding repetitive regions of the genome, and a relatively small percentage were found in the subtelomeric regions of the chromosome. In contrast, SNPs did not differ in distribution across the parasite chromosomes^45^.

We were therefore interested to see if *PfRad51* or the TLS polymerases played any role in this diversification process. We took the SNPs and indels data we obtained from the irradiated parasites and grouped them into those located in the core genome and those in the subtelomeric *var* clusters (Fig 4B and E). We observed that the three knockouts displayed a slight reduction (statistically significant in *PfRad51*Δ) in the number of SNPs in the core genome (Fig 4B). Surprisingly, the subtelomeric regions displayed an opposite phenotype where the SNP accumulation was slightly higher than in the wild type (Fig 4B). In contrast, the loss of *PfRad51* resulted in reduced indels in the core genome (Fig 4E), while the TLS knockouts did not show any significant changes. However, in the subtelomeric regions, the loss of *PfRad51* led to a significant increase in indels (Fig 4E). Our *PfRad51*Δ data showed the opposite of what we saw in 3D7 WT and published data. Sequence analysis from parasite clone trees from various *P. falciparum* isolates found that indels were concentrated in the noncoding repetitive regions of the core genome^2^. Our data implicate *PfRad51* in the repair of damage in repetitive regions in the core genome only. We could not confidently determine if the same scenario was prevalent in the internal *var* clusters due to the low incidence of SNPs and indels. Of note, the indels analyzed in this experiment are < 5 bases long, and though they could be included in larger recombination events, we instead saw a decrease in large recombinations as detailed below.

### *PfRad51* loss abolishes radiation-induced translocations in *P. falciparum*, while TLS polymerases play a minor role in recombination events

We performed a more detailed analysis of recombination events within the subtelomeric regions and the effects of the loss of HR or TLS pathways. To this end, we analyzed genomic DNA isolated from the three times irradiated clones and performed nanopore sequencing as outlined in Fig 5A. Nanopore sequencing provides read lengths that can extend over entire recombination events, including those within clusters or tandem arrays of multicopy gene families, which was not possible with Illumina sequencing. We expected most translocations to occur at the *var* clusters since they are more prone to recombination^8^. Our analysis revealed that the WT 3D7 exhibits some translocations after irradiation (∼3-4 in a clone), with most translocations happening within the sub-telomeric regions (Fig 5B, Supplementary Fig 4), as expected^10^. We observed no verified translocations in a clone of irradiated *PfRad51*Δ (Fig 5B), consistent with the anticipated complete loss of recombination in the absence of HR. However, this *PfRad51*τι clone displayed a large deletion in the AT rich region of the EMP1-trafficking protein gene (Pf3D7_1002100, Supplementary Fig 5). This deletion in the core genome indicates that alternative Rad51-independent HR repair pathways are present in the parasite, leading to large deletions but not translocations. The loss of TLS polymerases showed a similar phenotype to the WT 3D7, with 3-4 translocations per clone. Our results also show an increased propensity of the subtelomeric *var* clusters to undergo DNA breakage and recombination as most of the recombination events in all the lines happened between two ends of chromosomes, thus facilitating the generation of new *var* gene combinations as previously described ^3,7–9,47^.

**Figure 5:**
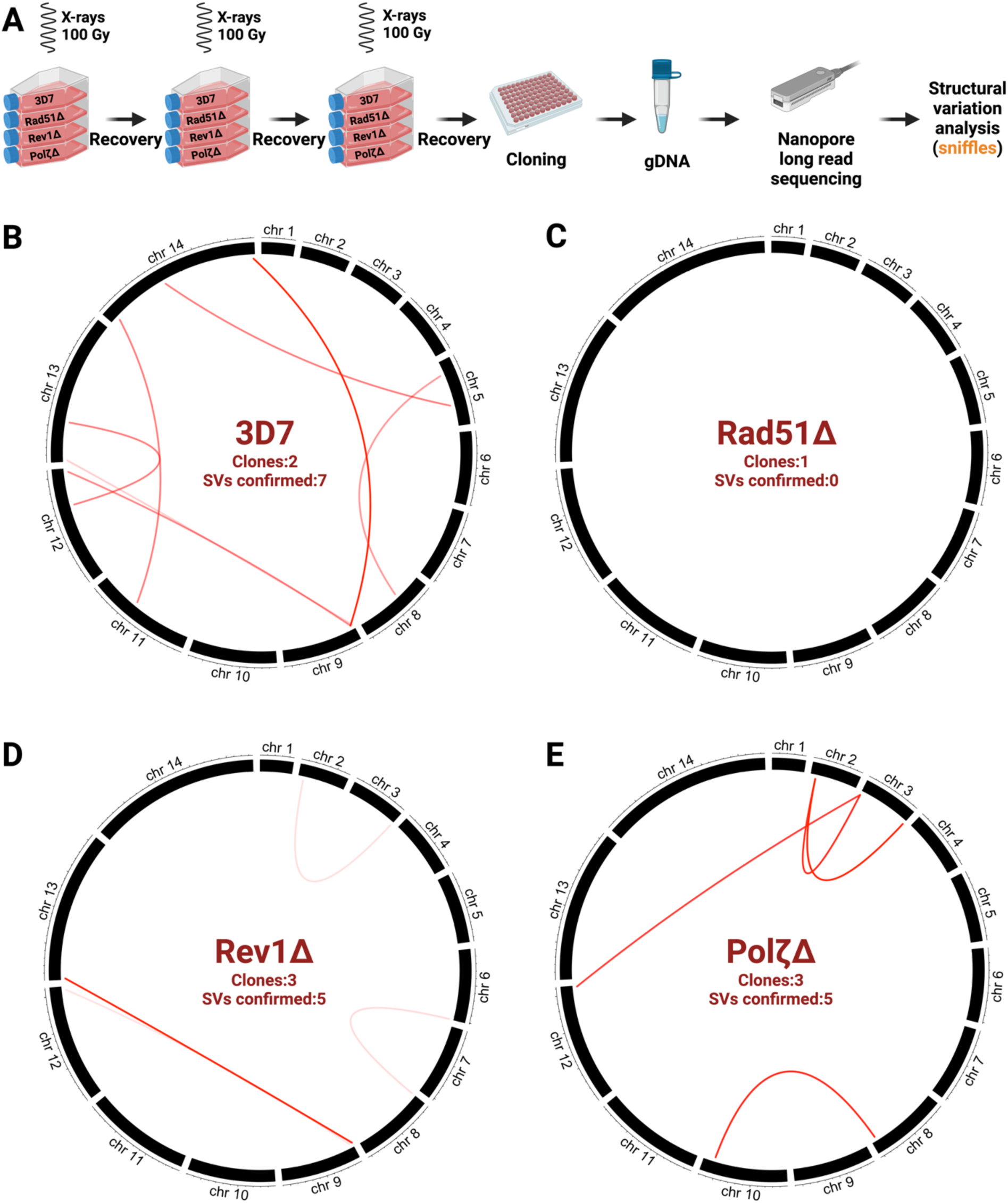
Irradiation induced translocations/recombination. (A) Overview of the experimental setup for the determination of recombination events. The translocations were confirmed by visual confirmation before plotting as described in methods. (B-E) Circos plots of translocations/recombinations distributed throughout the genome of *P. falciparum* cell lines – (B) WT-3D7, (C) *PfRad51*Δ, (D) *PfRev1*Δ, and (E) *PfPol*ζΔ after 3x irradiation with 100 Gy. Each line represents a recombination event between two genomic loci identified by the breakpoints. The visual IGV screenshots of some translocations in the WT-3D7 cell line have been described in supplementary Fig 3.

### *PfRad51* is essential for DSB repair in *P. falciparum*’s core genome, while TLS polymerases are dispensable

Based on previous reports and our results showing differences in DNA repair between core and subtelomeric regions (Fig 4F), we tested repair dynamics at the core genome and subtelomeric regions^4^. Our results above were generated from X-ray irradiated parasites where damage was global. Therefore, to study the effects of the loss of *PfRad51* and PfTLS polymerases on DSB repair dynamics at specific loci in the genome, we designed a CRISPR/Cas9-based DSB repair experiment. We targeted the gene PF3D7_1005500, which codes for *PfUpf1*, a component of the nonsense-mediated decay pathway in *P. falciparum* that is non-essential^48^. In addition, we determined that the knockout of *PfUpf1* does not affect parasite growth or any of the DNA repair mechanisms in *P. falciparum* ^49^. We added a variable in the distance from our directed cut to the homology blocks provided for repair by creating two plasmids containing repair templates for the *PfUpf1* gene, one with homology blocks that are ∼425 bp apart from each other in terms of their location in the genomic DNA, which we called “p*Upf1*Δ” (Fig 6A) and the other with homology blocks that are ∼3 kb apart from each other, called “p*Upf1*Δ-distant” (Fig 6C). The donor plasmids also supplied the single guide targeting the *PfUpf1* gene, as depicted in Figs 6A and 6C. Each plasmid was transfected into WT 3D7 and the knockout cell lines - *PfRad51*Δ, *PfRev1*Δ, and *PfPol*ζΔ, with a Cas9 expressing plasmid. Transfections were selected for 4 days with the required drugs for the Cas9 and donor plasmids. Each transfection was repeated 3 times.

**Figure 6:**
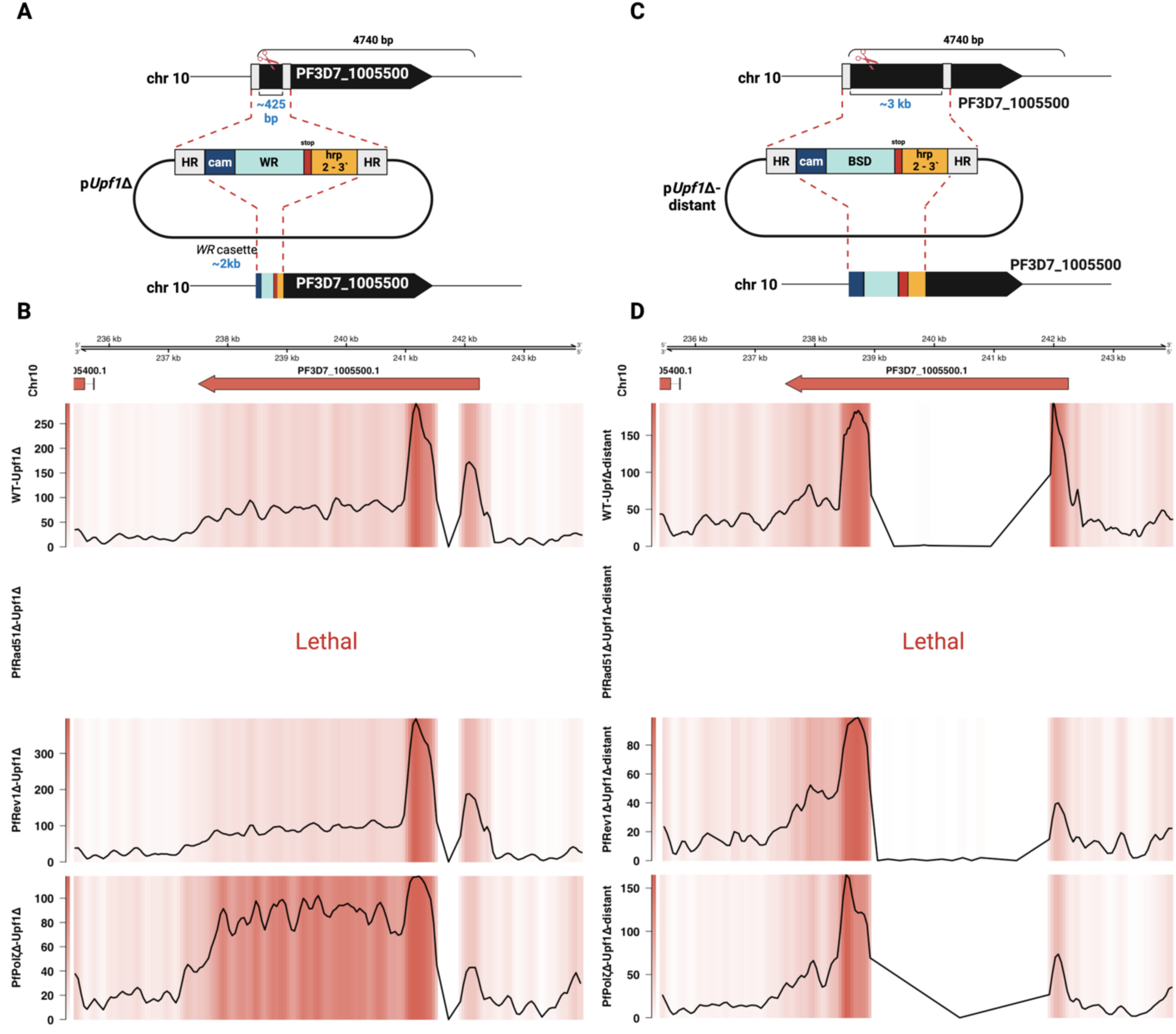
CRISPR/Cas9 mediated cuts to study homologous recombination in core genome of *P. falciparum*. (A and C) Overview of construction of *PfUpf1*Δ and *PfUpf1*Δ-distant plasmid constructs. (B and D) Representative coverage plots of Nanopore/Illumina sequencing results of *PfUpf1*Δ and *PfUpf1*Δ-distant in WT-3D7, *PfRad51*Δ, *PfRev1*Δ, and *PfPol*ζΔ. The top panel denotes the genome region, chromosome and the gene. The bottom panel has both heatmap and lineplot to show the integration site where the hDHFR/Blasticidin resistance cassette has been integrated which can be seen as a drastic drop in the coverage (Large dip in the lineplot coinciding with the white region in the coverage plot).

In WT parasites and both TLS knockout lines, we observed successful and efficient DSB repair by HR (Fig 6B). Specifically, in the WT, *PfRev1*Δ, and *PfPol*ζΔ strains, we recovered parasites with the predicted double cross-over event, resulting in the replacement of the sequence targeted by both the p*Upf1*Δ and p*Upf1*Δ-distant constructs (Fig 6B, 6D). This was expected as we are targeting a dispensable housekeeping gene, and as *PfRad51* is intact in all these lines, HR pathway should be functional. However, in the absence of *PfRad51*, the Cas9 induced double-strand break (DSB) proved lethal, with both donor plasmids, the p*Upf1*Δ and p*Upf1*Δ-distant, unable to serve as homology for repair in the *PfRad51*Δ cell line. Thus, *PfRad51* is essential for the repair of DSBs in the core genome, and TLS polymerases have a limited or no role in standard HR in *P. falciparum*.

### A *PfRad51*-independent homologous recombination pathway operates in *P. falciparum* subtelomeric regions

To study repair dynamics within the subtelomeric regions, we generated two repair plasmid constructs targeting the *var2csa* locus near the end of chromosome 12: *var2csa*Δ and *var2csa*Δ-distant, similar to the donor plasmids made to study DSB repair in the core genome and shown in Fig 7A and C. We chose *var2csa* as it is more conserved between *P. falciparum* isolates and distinct enough from other *var* genes to reduce the likelihood of ectopic recombination events with other *var* genes^50,51^. The *var2csa*Δ and *var2csa*Δ-distant constructs contained repair templates ∼1.5 kb and ∼2.5 kb apart in the genome, respectively. In addition, the *var2csa*Δ-distant plasmid had no selection marker between the two homology blocks as opposed to the *var2csa*Δ plasmid (Fig 7, A and C). Similar to the core genome experiments, each plasmid was transfected into the WT 3D7 and the knockout cell lines - *PfRad51*Δ, *PfRev1*Δ, and *PfPol*ζΔ, with a Cas9 expressing plasmid. We selected for the repair plasmid for 4 days after transfection. Again, repair plasmids also supplied the single guide targeting the *var2csa* gene.

**Figure 7:**
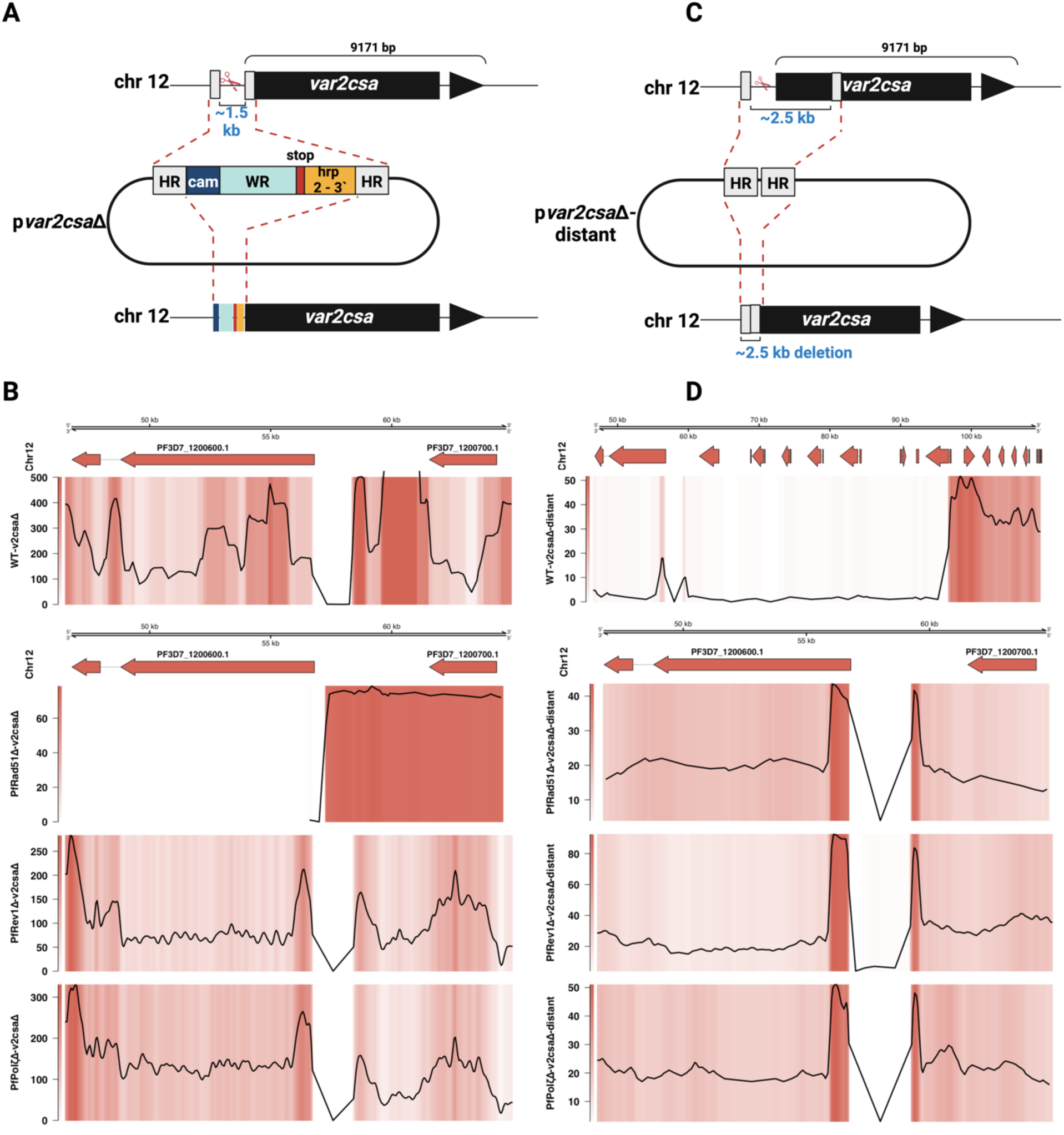
CRISPR/Cas9 mediated cuts to study homologous recombination in subtelomeric regions of *P. falciparum*. (A and C) Overview of construction of *Pfvar2csa*Δ and *Pfvar2csa*Δ-distant plasmid constructs. (B and D) Representative coverage plots of Nanopore/Illumina sequencing results of *Pfvar2csa*Δ and *Pfvar2csa*Δ-distant in WT-3D7, *PfRad51*Δ, *PfRev1*Δ, and *PfPol*ζΔ. Note that var2csa is abbreviated as v2csa in the figures. The top panel denotes the genome region, chromosome and gene. The bottom panel has both heatmap and lineplot to show the integration site where the hDHFR cassette has been integrated or the ∼2.5kb DNA piece (as shown in C) has been lost due to HR, which can be seen as a drastic drop in the coverage (Large dip in the lineplot coinciding with the white region in the coverage plot). Please note that in D, the WT-*Pfvar2csa*Δ-distant sequencing data shows a larger genomic region compared to the other samples to efficiently cover the large resection of ∼35kb leading to TH.

In subtelomeric regions, DSBs can be repaired either by *PfRad51* mediated HR or by telomere healing (TH) ^8,10,37^, a process in which the fragment between the site of the break and the chromosome end is lost, and a telomere is added de novo to the DSB site. Parasites can survive TH as the subtelomeric regions are not essential^8,10,37^. In the WT background, the *var2csa*Δ construct resulted in efficient HR in which the two homology blocks were utilized to repair the DSB (Fig 7B). In contrast, the loss of *PfRad51* abolished HR and forced the parasites to repair the DSB through TH (Fig 7B). As observed in the core genome, the two TLS polymerase knockouts phenocopied the WT, indicating that the *PfRad51-mediated* HR is functional at all chromosome locations in these knockouts (Fig 7B).

Surprisingly, in the WT background, when the *var2csa*Δ-distant construct was provided as the template for repair, we observed that the repair machinery failed to recognize the homology blocks, resulting in a large (∼35kb) resection and TH (Fig 7D). This result indicates that despite a functional HR pathway within the subtelomeric regions, TH outcompetes HR when there is a significant distance between homology blocks. In contrast, TH is not an option in the core genome, and therefore, HR proceeded as expected even when the distance between homology blocks was increased (Fig 6D). More surprisingly, we observed homology-mediated repair of a subtelomeric DSB in the *PfRad51*Δ line despite the loss of the HR pathway. This was unexpected and also observed in the two TLS knockouts (Fig 7D) indicating that when the distance between the two homology blocks was greater than 2 kb, in the absence of *PfRad51* or the two TLS polymerases, we saw the activation of an alternate Rad51-independent pathway that can perform homology-directed repair specifically within the subtelomeric region. In the absence of Rad51 or the TLS polymerases DSB repair was skewed towards a different result than what was seen in the WT 3D7. When we examined the individual reads that spanned the DSB site in all three knockouts, we observed a recombination mechanism that identified the first homology block but initially failed to recognize the second homology block and instead replicated the entire plasmid backbone multiple times before using the second homology block to complete recombination (Supplementary Fig 6). This resulted in the incorporation of multiple copies of the plasmid at the site of repair in all three of the knockout lines when our DSB targeted a subtelomeric location. The same result was seen in multiple repeated transfection experiments (3 transfections each). Previously, others have described similar integration events in non-CRISPR mediated genetic modifications in Plasmodium. It is distinct, however, that with the same guide and donor plasmid, we noted very different repair events in our WT versus knockout parasite lines. Notably, we did not observe this type of repair within the core genome.

## Discussion

We set out to identify mechanisms of malaria genome diversification with a particular interest in how DNA repair mechanisms would differ at different chromosome locations in *P. falciparum* and are now just beginning to understand how the parasite segregates its genome into a well conserved core genome and highly divergent subtelomeric regions^4^. We previously proposed that interactions between the TLS pathway and HR in malaria were critical to diversifying the parasite genome, and now, in this current work, we have pursued the mechanism behind these interactions and parasite genome diversification^17^.

*var* gene diversity has been described by our lab and others, ^9,47,52–54^ and recent work is starting to address the possible mechanisms that underlie *var* gene diversification ^7,8,17^. Analysis of recombination events showed that, in general, *var* gene recombination maintained open reading frames and preserved the domain architecture, resulting in functional chimeric *var* genes. This is consistent with gene conversion and NAHR as the primary mechanisms to generate novel *var* gene sequences during mitotic replication^7^. This idea is further supported by work describing recombination identified in parasites subjected to irradiation and targeted DSBs via CRISPR/Cas9^8,10^. Both modalities of DNA damage yielded subtelomeric gene recombination consistent with a cascade of ectopic HR events leading to the formation of chimeric *var* genes^8,10^. To date, these ectopic HR events have been primarily observed in the subtelomeric *var* clusters where the repair choice between TH and HR drives diversification^10^. We have yet to understand alternative diversification events that occur amongst the five *var* gene clusters located internally on chromosomes 4, 7, 8, and 12. HR is historically considered a non-diversifying DNA repair pathway that maintains the overall gene structure, and we propose that this very characteristic is adapted by the parasite to recombine and produce chimeric *var* genes by using the redundancy in *var* gene sequences.

One of the most striking observations is that *PfRad51*, *PfRev1*, and *PfPol*ζ are not essential for the survival of parasites and that there was no growth defect under standard culture conditions, even given the replication stress the parasite is under during erythrocytic development as it replicates from one ring to 30-50 infectious merozoites (Supplementary Fig 2). This contrasts with the essentiality of Rad51 in vertebrates^23^ but aligns more closely with observations in lower eukaryotes such as yeast ^24,26^. The stage-specific differential sensitivity to irradiation between the ring and late stages is primarily seen in the TLS knockout parasite lines and reveals stage-specific DNA repair mechanisms in *P. falciparum* similar to dominance of different DNA repair pathways at different stages of the cell cycle in higher eukaryotes^55^ (Fig 2A, 0-24 hours post-invasion - hpi, and Fig 2C). Particularly noteworthy is the essential role of TLS polymerases only in ring-stage parasites’ survival following irradiation (Fig 2C). Ring stage parasites are haploid and are not undergoing DNA replication and, therefore can be considered analogous to the G1 phase in higher eukaryotes such as fission yeast^56^. In the late stages (Fig 2A, ∼24-36 hpi), which could be compared to the S-phase of higher eukaryotes ^56^, HR appears to dominate repair due to the availability of a second copy of the DNA, and in this setting, the TLS polymerases are dispensable. The differences in stage-specific requirements of TLS polymerases highlight the complex and dynamic nature of DNA repair in the parasite.

Since *P. falciparum* lacks the error-prone NHEJ pathway, we predicted that the subsequent loss of *PfRad51* would result in a large majority of DNA breaks being unrepaired and thus lethal. The slightly longer recovery for later stage *PfRad51*Δ parasites was not as pronounced as expected and suggests that PfRad51 may have a non-HR related role in the ring stage parasite response to DNA damage that is worth exploring in future work. Analysis of whole genome sequencing (WGS) data or our irradiated knockout parasite did not highlight the dominance of an alternative DSB repair pathway to step in for HR. We found one example of a large deletion between repetitive regions consistent with a Rad51 independent HR termed Synthesis Dependent Strand Annealing (SDSA) which is characterized by deletions within a repetitive region and mediated by Rad52. However, to date, no Rad52 ortholog has been identified encoded in the parasite genome. Nor did we see an increase in insertions or deletions consistent with MMEJ, which has been described in *P. falciparum* but appears to be limited even in the setting of no C-NHEJ or HR (*PfRad51*Δ).

Irradiation results in large-scale genome damage that can lead to the accumulation of mutations in the form of SNPs, indels, and translocations. The decrease in SNPs in the TLSΔ lines was expected and is consistent with TLS polymerases’ central role in the DNA damage response and knockouts leading to a hypomutable state. For Plasmodium, the increase in SNPs in our data suggests a more error prone alternative pathway is available to the parasite in the absence of *PfRad51*. An increase in SNPs in a Rad51Δ has been noted in other organisms such as Leishmania^57^. Interestingly, in engineered parasite lines missing the editing function of Polδ (created in both *P. falciparum* and the rodent malaria *P. berghei),* elevated mutation rates were seen and were increased specifically in coding regions^45,46^. In addition, the mutation rates reported were significantly higher in the *P. berghei* Polδ mutant. All rodent parasites lack the TLS pathway, which may have a role in the differential mutation rate between Plasmodium species^17^. When we exposed parasites to DNA damage in our system, the TLS Δ parasites could not survive and were, in contrast, hypomutable.

Genome analysis of parasites grown in standard culture conditions and then subcloned identified similar mutation rates between *P. falciparum* isolates of different geographical origins and an even distribution of SNPs across coding and noncoding regions^2,7,9^. This work also noted a predominance of new indels in noncoding AT-rich repetitive sequences in the parasite core genome^2^. The authors proposed that the sequence of exons and noncoding regions were associated with distinct forms of indel mutations. Our data show that the loss of *PfRad51* resulted in a significant reduction in the number of indels in the core genome, and thus, Rad51 is likely to play a role in indel formation. Interestingly, we did find that the subtelomeric regions had an increase in the number of indels in *PfRad51*Δ compared to the WT (Fig 4E), the opposite of what is seen in WT parasites^2^. This difference suggests that different DNA repair proteins may respond to damage at different chromosomal locations, perhaps due to epigenetic or other factors that might maintain a demarcation between core vs subtelomeric regions of the parasite chromosomes.

Our results show that *PfRad51* is important for structural variations (Fig 5B). We proposed that increased ectopic recombination occurring specifically in the subtelomeric regions plays an important role in this diversification ^8,10^. This is also supported by others who observed that while SNPs were scattered across the genome, SVs were focused in and around *var* genes^7^. Hamilton et al. also found that while the SNP rate did not differ between parasite isolates, the frequency of *var* gene recombination through NAHR did, with more recent clinical isolates demonstrating a higher rate of chimeric *var* gene formation^2^. Potential influences, such as *var* gene transcription, remain to be studied. We utilized CRISPR/Cas9 directed cuts at a specific locus in the core genome and one of the *var* clusters in the subtelomeric region to determine the repair dynamics and how the loss of *PfRad51* and the two TLS polymerases affected DSB repair dynamics (Fig 6 and 7). Our results show a clear difference in the choice of DNA repair mechanisms depending on whether the break occurs within the core genome or a subtelomeric region. In subtelomeric regions, parasites can either repair a DSB via HR or stabilize the chromosome end by TH, consistent with our results. In addition, the free fragment formed after the DSB within a subtelomeric region has been shown to be highly recombinogenic and lead to ectopic recombinations readily ^58^. This would also explain why the multicopy gene families are clustered in the subtelomeric regions in apicomplexans, thus enabling a higher degree of spontaneous diversification.

Further, our experiments with the distant construct, which we expected to function as a ‘non-perfect’ repair template, showed a clear differential DNA repair between core vs subtelomeric region (Fig 6D and 7D). While *PfRad51* is essential for repairing DSBs in the core genome (Fig 6B), it is not essential in subtelomeric regions (Fig 7B) as the parasites can choose TH over HR. However, in the subtelomeric region when the distant repair template was provided, the choice of repair was HR, and to our surprise, this HR was *PfRad51* independent (Fig 7D). This difference between core and subtelomeric regions might contribute to the highly diverse nature of the subtelomeric regions. The question that remains unanswered is, how do parasites perform HR, specifically in the subtelomeric regions in the absence of Rad51. Further exploration into proteins that play a role in this process has the potential to identify novel mechanisms that enable human malaria parasites to diversify the *var* repertoire and thereby evade the human immune response more efficiently by switching to novel, chimeric *var* genes.

The *P. falciparum* genome is unique in many aspects and can serve as a model for the study of DNA repair under high oxidative and high replicative stress in a haploid organism that lacks NHEJ. The evolution and regulation of DNA repair pathways in the malaria parasite can serve as guides not just to antigenic variation and drug resistance in this parasite of global importance but also to DNA repair mechanisms in other eukaryotes.

## Materials and Methods

### Culture and genetic manipulation of parasites

Parasites were cultured in a standard culture system at 5% hematocrit in media containing RPMI 1640 (Corning Life Sciences, Tewksbury, MA, USA), 0.5% Albumax II (Invitrogen, Carlsbad, CA, USA), 0.25% sodium bicarbonate, and 0.1 mg/ml gentamicin in an atmosphere of 5% oxygen, 5% carbon dioxide, and 90% nitrogen. Culture media was supplemented with the required selections (WR, Blasticidin (BSD), or DSM1) as per the genotype of the cell lines. Clonal parasite lines were obtained by limiting dilution ^59^. Parasites were transfected by electroporation^60^ using derivatives of the plasmids pL6_eGFP and pUF1_Cas9 for CRISPR/Cas9-based genome editing as described by Ghorbal and colleagues^28^. Homology blocks for genome modification were amplified from parasite genomic DNA by PCR, single guide sequences were synthesized as oligos, annealed, and both were inserted into pL6 by infusion cloning (Clontech, Takara Bio USA, Mountain View, CA, USA). The plasmids generated are listed in Supplementary Table 1. Single guide sequences and sequences used as homology blocks for CRISPR modifications are listed in Supplementary Tables 2 and 3. Growth of the parasites was monitored using flow cytometry as described before^61,62^. The parasite cell lines used in this study are listed in Supplementary Table 4.

### Genomic DNA extraction

Infected red blood cells from a 10-ml culture (∼4-5% late stages) were centrifuged at 4,000 rpm. The supernatant was discarded, and the pellet was resuspended in 1 ml of phosphate-buffered saline and 15 μl 10% saponin. The parasite pellets were spun at 12,000 rpm, 3 mins, and washed twice in 1 ml of PBS. The pellet was then taken up in 400 μl Tris-sodium chloride-EDTA buffer, along with 80 μl of 10% SDS and 40 μl of 6 M NaClO4. This suspension was placed on a rocker at room temperature overnight. The following morning, an equal volume of phenol-chloroform-isoamyl alcohol (25:24:1) was added and gently vortexed (for Nanopore sequencing samples vortexing was skipped to maintain the length of the DNA and gently mixed by inverting the tube 5-10 times) and centrifuged at 16,000g for 5 min. The final aqueous phase was ethanol precipitated and resuspended in 50 μl of sterile distilled H2O. The final DNA concentration was determined by absorbance at 260 nm using a nanodrop machine.

### Sequencing of parasite genomic DNA

Parasite genomic DNA was extracted using phenol-chloroform extraction, and sequencing libraries were prepared from 1 μg of unsheared gDNA (as measured using the Qubit dsDNA BR assay [Life Technologies, Carlsbad, CA, USA]) using the Oxford Nanopore Technologies ligation sequencing kit version 1D LSK 108 (Oxford, UK). This approach minimizes the potential shearing of DNA template fragments and entirely avoids PCR artifacts by ligation of a manufacturer-supplied adapter nucleoprotein complex to native gDNA to facilitate the loading of long library molecules into the protein nanopores. Nanopore sequencing was conducted with an Oxford Nanopore Technologies GridION X5 instrument using MIN-106/R9.4.1 flowcells. All samples were sequenced using a standard 48 hr run controlled by Oxford Nanopore Technologies MinKnow software, and raw data were basecalled with Albacore version 2.2.4. FASTQ files were extracted from basecalled FAST5 files with Poretools ^63^.

Illumina whole genome sequencing was performed using NovaSeq 6000 system to get about 5 million reads per sample. Sample library was prepared using Kapa Hyper library prep kit (Roche) and run on an S2 flow cell with 2×100 cycles to obtain paired-end reads in the fastq format. All the sequencing data can be accessed at: https://www.ncbi.nlm.nih.gov/bioproject/PRJNA1244413

### Irradiation of parasites

Irradiation of the parasites was performed using RS 2000 Biological Research X-ray Irradiator (Rad Source Technologies). Briefly, parasite cultures were exposed to the required amount of irradiation utilizing the level 3 shelf, which supplied ∼2.1 Gy/min. The culture was maintained by media change and by replenishing blood until the parasites that survived the irradiation came up. Once up, the parasites were irradiated again for the second time and repeated for a third time. After the third round of irradiation, the gDNA from the surviving parasites were isolated as mentioned above using the PCIA extraction, purified using Zymoresearch genomic DNA clean and concentrator kit (Cat.no - D4010), and Illumina sequenced as above for quantifying SNPs, Indels or used for determining *var* gene copy number variations using qPCR.

### Quantification of SNPs, Indels, and translocations

To quantify SNPs and Indels from the Illumina sequencing datasets, reads were trimmed using Trimmomatic ^64^ to remove adapters, read quality was checked using FastQC^65^, and the genome was indexed and aligned using BWA-MEM^66^. Duplicates were marked using sambamba ^67^ and the bam file was sorted and indexed using samtools^68^. Variants were called using freebayes^38^ and the vcftools package was used to filter out variants having quality (QUAL) scores less than 30 and DP scores less than 10. To determine the translocations from nanopore reads, bam files from the raw fastq files were generated using minimap2^69^, sorted and indexed using samtools^68^, and variants called using sniffles^70^. SNPs, Indels, and translocations unique to the irradiated cell lines were identified using the vcf-isec command of vcftools ^71^ and were plotted into circos plots using the R package circlize^72^. Translocations identified by sniffles were visually confirmed using IGV^73^ before circos plotting. All the codes for the analysis performed can be accessed at GitHub here: https://github.com/agnibhat/Characterization-of-Plasmodium-falciparum-DNA-repair-dynamics.

## Acknowledgements

We would like to thank Björn F.C. Kafsack for insightful inputs on the project and experiment design. Richard McCulloch for helpful discussions. We would also like to thank the Genomics core at Weill Cornell Medicine for the Illumina sequencing services.

## Conflict of Interest

None declared

## Funding

This research was funded by the National Institutes of Health, grant number R01AI146153. The content is the responsibility of the authors and does not necessarily represent the official views of the NIH.

## Data availability statement

All the sequencing data have been deposited to the SRA database (https://www.ncbi.nlm.nih.gov/bioproject/PRJNA1244413). All the codes for the WGS analysis performed is available at GitHub (https://github.com/agnibhat/Characterization-of-Plasmodium-falciparum-DNA-repair-dynamics)

## Notes

https://www.ncbi.nlm.nih.gov/bioproject/PRJNA1244413

### Competing Interest Statement

The authors have declared no competing interest.

### Summary of Updates

Acknowledgements: We would like to thank Bjorn F.C. Kafsack for insightful inputs on the project and experiment design. Richard McCulloch for helpful discussions. We would also like to thank the Genomics core at Weill Cornell Medicine for the Illumina sequencing services. Conflict of Interest: None declared Funding: This research was funded by the National Institutes of Health, grant number R01AI146153. The content is the responsibility of the authors and does not necessarily represent the official views of the NIH.

