## Supplemental files for "*Plasmodium falciparum* DNA repair dynamics reveal unique roles for TLS polymerases and PfRad51 in genome diversification"

### Supplementary Data

Supplementary Table 1: List of plasmids generated from pL6-eGFP as described in the methods section.

| Plasmid name | Description |
|---|---|
| pRad51Δ | Generating knockout of *PfRad51* |
| pRev1Δ | Generating knockout of *PfRev1* |
| pPolζΔ | Generating knockout of *PfPolζ* |
| pUpf1Δ | Generating knockout of *PfUpf1* |
| pUpf1Δ-distant | Generating distant version knockout of *PfUpf1* |
| pvar2csaΔ | Generating knockout of *Pfvar2csa* |
| pvar2csaΔ-distant | Generating distant version knockout of *Pfvar2csa* |

Supplementary table 2: DNA sequences used in the study

| **Oligo Sequence** | **Target** |
|---|---|
| TTATTTGGTGAATTTCGTAC | single guide for generating *PfRad51*Δ |
| ATATTAATAGTTCAAAACGG | single guide for generating *PfRev1*Δ |
| TTATCTTTTTAATGAAAGTG | single guide for generating *PfPolζ*Δ |
| TGAATGCAATCATTACCATG | single guide for generating *PfUpf1*Δ |
| TGGAATAAAGAAACATACTT | single guide for targeting *var2csa* |

Supplementary table 3: DNA sequence of homology blocks used for generation of cell lines.

***PfRad51*Δ**

Homology block 1:

CTGAAACAAGCAAATACAAAAGAAGATAAGTCGCAGAAAATTTCCAATAGTAGTAC
TATTGATGAAATAGAAGAAGAACAGTTATATACTGGACCTTTAAAAATAGAACAAT
TATTAGCAAAAGGTTTTGTAAAAAGAGATTTAGAATTACTGAAGGAGGGTGGATTA
CAAACTGTTGAATGTGTGGCATATGCTCCTATGCGTACGTTATTGCTATTAAAGGTA
TAAGTGAACAGAAGCAGAGAAATTAAAAAAGGCAGTAAAGAATTATGTAATTCAGG
TTTTTGTAATGCTATAGATTATCACGACGCAAGACAAAATTTAATAAAATTTACCAC
AGGATCAAAACAATTAGATGCCC

Homology block 2:

CTATTACATGTCAATTACCTATTGAACAATCAGGAGGAGAAGGTAAATGTTTATGGA
TAGATACGGAAGGTACTTTTAGACCTGAGCGTATTGTAGCTATCGCTAAAAGGTATG

GTTTACATCCAACGGATTGTTTAAATAATATAGCATATGCTAAAGCCTATAATTGTG
ATCATCAAACTGAATTATTAATAGATGCTAGCGCTATGATGGCAGATGCAAGATTTG
CTTTATTAATTGTGGATTCAGCAACGGCTCTATATAGATCTGAATATATAGGTCGAG
GTGAGCTAGCTAATAGGCAATCACATTTATGTCGA

***PfRev1*Δ**
Homology block 1:
GATTCAGTGGAGGATTCAAAAGAATTAAACAATTATATGAATAACAATAGTATTAA
TAAAACAACGTTAGGTAAGTTTAATATGGAAAATAAAATGGAATATAATTATACTTC
TATTAACGAACATAATAATAATAATTATTATATAAATAAGATACATAATGATTCTGT
TTTTAAATATCAAGATATGGATAAAACACCTATGAAAAATAACATAGATAATAATA
GTAATAATAATAATAATAATAATGATAATAATAATAATAATGGTAATACATATTTGT
ATGATAATTATCTAGAAGACCAATTTAATCAAACAATAAAAAATCGATATAGTTGTA
CTCCCAATGATCAAATAAGCACTTGTACAAAAAGTTTTAATACATGTATAAAAACAT
ATGAAGAACAAGCTTCTTCAAGTTGTAAAAA

Homology block 2:
GACCAGCAAAGTAACACACATTATTAGTAATAACATGGCATTAGGTTCTAAGAAAT
ATATGGATTATAAAAAAGCTATAAAAAAATCTAAAGTCTTCATAGTTATAGATCAAT
ATATTTTCGATTGTGTAAACATGCAATGTCGTTTACCTGAACAATCATATCTACCTTC
CATGTTACGTTACAATTGTCATCAAATAACCGAATATTTTTCCTTAAGAAAAAAAGA
TAAGGAACAAAACAAAAAAATGCAAAACAAAAAAATACAAGACAAAATAAATAAT
GATCAAAATTTTTTGAAGGAACATAAAGAAGGTG

***PfPolζ*Δ**
Homology block 1:
TGAGAAAGGATGCATTGCATCATTCGAAATTAATAAAAAGAGAAAAGATGTGGTTC
GTTATTCTTTTAAGAAGGAACCACCTAGAATAAACAAATGTTATGCCGTTATTAATA
ATTTAGTTGATCATATAAGTGGAAGAGATAACAAAAATATGAAAGAAAAAATGAAA
GGAAATATTTATAATAGAATACATGACAATATAGAAGATGAGAATAAAGAAGATAC
ATCAAAATTTGAATATATAGGAAAAGAAAATCATATGGAAAATAAGGAAAATATAA
GAAAGCAATATGAACAGATCAAATCAGATAATATGAAGAAAAAAATAAATATAAA
ATATGGAAATATATTTTTTCTAGAAATATTAACCGAAATAAAAGATGAAAATTGTTA
TTCTTCAGATTATAACCAAGACAAAATAAAGGCCGTTTTTATATAGTGAGGGAGGA
AAGACTTATG

Homology block 2:
GGAACAAATGAAGTCAAATTAAATGAATCCAAATTAAATGAATCCAAATTAAATGA
ATCCAAATTAAATGAAGAAGAATTAAAATTGTGTAAAGATATGTATCATATAGAAG
ATGTGAACCTTTGTACAAATAATTGTAGAAGTATAGAAAAAAATAGGGATAATCTA

ATAAATGATAAGAATGTGATTAATAAAAAGAACGACGATACATATGGAAGTAGTAA
AGAATTATGTTGTAGTAAAAATGGTTATCATACAAATAAAATAATAAAAGAAATAA
CAAATAATGATATAAAAAAGGTGAAGAGAACGTTTTTGATTTTAATATAAATTATA
ATAACGTTAATATATGTATTGTAGAAAATGAAAGAGAACTTATTCAAAAGTTGATTA
ATAAGATATTGTTTTATTCTCCGCTTAGTATTGTTTCTTATGAAAATGATAAATATAA
TATAAATTATATAAACCAAAGATGTTTAGCTTTAGATATAGG

***PfUpf1*Δ**

Homology block 1:
CATTATCCACGTCGTGATCTTCTAACAAAATCTCCTTTTCCATATCATCTTGTAAACT
CTTACTTTTGTGTTCATCAAAAGGACCGACATGTTTTCATATAAGGATATATAACTA
GATAAACAAGGATCTCTACAAATAATAACAACAACACCTTCTTCGGATGTCGGAAG
GAAACCTAACAAAAATACATTTCTGCATGCACAGTTATAACATTCTAAAATTGTTTC
ACCTAATAAACTATTTTTATGTAATCTAATTTCTTTATGTTTTGATCGAACTAAATGT
GTAACAATGTGACTACCACAAGTACCATAAGAACCATTACAAAACCAACGCTTACA
ATTATTACATTGAACAACACTATCAATAGAATCAATTTCACAATATCTGCATCTATA
ATATTTTAAATCATCTTTACTTTTTTTATTTTTATATCTATAATTATTACCTTCTTCATA
AAGTTCATTATGTGTGTCATCCTT

Homology block 2:
ATTCCGTATCTTCCAAACTTTCATTTAACATATCTAACCTCTTTTCCTCTATTATTTTA
CATGGAGAATATTCCGATTTTTTATTTTTCCCATGTTTTTTTGTAGGTTTTATATTTGT
ATTTTTTTTTAACACCTCTTTATTACATGTATCCTTATCGTCATTAAATTTTTGTTGTG
TATCTTCTGAAAAATAGTTTTCTTCTTTTATTTTTTTTTTTATTCGTATCCTTAGAAATA
CAAAAATCGTACAGTTCGTCAAAATTATCTACATGAATATTATTATAATCGAAAGTT
TTATCCATAGTTTTTCATACGCCCCTACAGATGCTTTTATAAATAAATAAATAAATAT
ATATATATATATATATATATATATTATATATATATATAATTATATTTACTCAATTATA
TAAATAACCATTACAGCTATATAAAACTAATTTCTTTTTCTCCTTTTATTTTGTATGGT
CTAACATATAAGGTGTCCTATACACACTCCA

Homology block 3 (distant):
GGTTTCAAGTTAGCTAAACAACCTTCTACAATTAAATCTTTTTTTTTAAATTGACTTA
ATAAATTGATCCATACTGAATTTACATTAGTTATGGTTTCATTTGCATTAATTTTTTCT
CTGGATATAAAATGATGTCTTGATAAAACTTTTGCATTTCCACATATAATTAATCCAT
ATTTTGCTCTGGTTAAGGCAACATTTAATCTTCTAGGATCATTTAAAAACCCTATACC
TAATTTTTTATTAGATCGAACACATGATAATAATATAAAATCTTTTTCTCTTCCTTGG
AATGCATCTACTGAAGCTACTTCTATATCCGACGAATTTTGAAAAGAAATATTTTTTT
GAAATAATGAAGTAATATATGCTCTTTGTCCTTCATAGGGTGTTATAACACCTATTTG
TGATGGTTTCAAACCACATTGTAATAAGGTACGAACTAATTTTTCCATATTAGAAGC
TTCACTTCTATTAAGATAACTTGTACCTGATGCA

***Pfvar2csa*Δ**

Homology block 1:

ACTATAGAATACTCGCGGCCGCCATGAACGCTTAAAGAAACAAGGAAAAAGGAATA
TAATTACAAAACACTTATACACACATATATATATATATATATATATATATATATAT
GTATATATATACATATTATGGTAGTATTATTATAGGGTATAACTTTAATTTATATAAT
TTTATAATATATATAATTCCATAACTTTAACTAATAATATAATATTATTTATAATAAA
AATATCTGAAAATAAAGTACAAACATATTATAAATATAACTAAAGTATTATATATTT
ATAACAAATATAGATATAAAAAAATATTTGTATGAAATTCATTAAACTATATTTATA
ACATAACAGTGTTGTAATATATATAAAATATACAAATTATGTAACACCAAATATATG
TAAAAAAAAAAATATATTTATAAGAATATATCAAATTTATTTGACAACAAATTGTAA
AAATATAAAATTATAATAATATATTATTAAAATAACAGTAGTGAAATGATATTATAT
GTTGTTTATAAGATAAAGAAAATATTTTAATTATATAACATAAAGATAATATTCTTT
AAATATAATAATAATAATATTATAAATTTTAAATATAAAATTATTAAGTTTTTATGTA
AACTTAAAATATAATAAATAATTATATTATAATATATTTAGAAAAAATATATTTTTA
TAAAAGTATAATTATTTATTTTGACTTTTGTTTTTAATTGAACTCATTTAAATATCTAT
ATATATATATATATTCAATTATAATTATATATATGAAATATATAAAACTACTTAAAA
TATAATAAAAATAAAAACATGCCATATTAAAAAGAAGAAATTATATTTTAAATTATA
ATAATTATTATAATAACATTCTAAAATAATAAAGGTATATTTATAAGTTAAAAAGAA
TAAAAATTATGTATAAAATAAAATTATAACTATATACATATTATAAATAAGAATAGC
ATATACTAGTTTTTATTATACGACATAAAATTATATTTAGATTTAATGGAATACATAT
AGTACATAATAGTACATACATATATATATATTTTTTAATTGATATTTAACCATATATA
TATAAGATATTCAATTATAGTAAATGTTTTTATGTTTATTTATTTTATAGAGTGTATA
GATAGAAATGGAATACTTTATTTGCTTCCTT

Homology block 2:

GAATTCGACCAATCGTTGAAAGCTGATCCTAGTGAAGTGCAGTACTATGGAAGTGG
AGGTGATGGATATTACTTAAGAAAAAATATTTGCAAAATTACCGTGAATCATTCAGA
TTCTGGAACAAATGATCCTTGTGATAGAATACCACCTCCTTATGGCGATAATGACCA
ATGGAAATGTGCCATAATTTTATCTAAAGTAAGTGAAAAACCTGAAAATGTATTTGT
TCCTCCGAGAAGACAACGTATGTGCATTAACAATTTAGAAAAATTAAATGTTGATAA
AATTAGGGATAAACATGCATTTTTGGCAGATGTATTACTTACGGCCAGAAATGAAGG
AGAAAGAATAGTGCAGAATCATCCAGATACAAATAGTTCCAATGTTTGTAATGCATT
AGAAAGAAGTTTTGCTGACATTGCAGATATTATTAGAGGTACAGATCTATGGAAAG
GTACTAATAGTAATTTAGAACAAAATTTAAAACAAATGTTTGCAAAAATACGAGAA
AACGACAAGGTACTTCAAGATAAATACCCAAAGGACCAAAATTATAGAAAATTACG
AGAAGATTGGTGGAATGCTAATAGACAAAAGGTGTGGGAAGTTATTACTTGTGGTG
CGCGAAGTAACGATTTACTCATAAAACGTGGATGGAGAACATCTGGAAAATCTAAT
GGAGACAATAAACTTGAATTGTGTCGCAAATGTGGCCATTATGAAGAAAAGGTTCC
TACCAAATTAGATTATGTCCCTCAATTCTTAAGGTGGCCATGG

Homology block 1 (distant):
ACTATAGAATACTCGCGGCCGCCATGAACGCTTAAAGAAACAAGGAAAAAGGAATA
TAATTACAAAACACTTATACACACATATATATATATATATATATATATATATATAT
GTATATATATACATATTATGGTAGTATTATTATAGGGTATAACTTTAATTTATATAAT
TTTATAATATATATATAATTCCATAACTTTAACTAATAATATAATATTATTTATAATAAA
AATATCTGAAAATAAAGTACAAACATATTATAAATATAACTAAAGTATTATATATTT
ATAACAAATATAGATATAAAAAAATATTTGTATG

Supplementary Table 4: List of cell lines and clones and type of sequencing done

| Cell line/Clones | Type of Sequencing | Ref |
|---|---|---|
| WT(3D7) | Illumina/PacBio | This study[1,2] |
| *PfRad51*Δ | Illumina/Nanopore | This study |
| *PfRev1*Δ | Illumina/Nanopore | This study |
| *PfPolζ*Δ | Illumina/Nanopore | This study |
| WT-3x-irradiated clone D5 | Illumina | This study |
| WT-3x-irradiated clone F12 | Illumina | This study |
| WT-3x-irradiated clone G2 | Illumina | This study |
| WT-3x-irradiated clone H10 | Illumina | This study |
| WT-3x-irradiated clone E8 | Illumina | This study |
| *PfRad51*Δ-3x-irradiated clone B11 | Illumina | This study |
| *PfRad51*Δ-3x-irradiated clone D11 | Illumina | This study |
| *PfRad51*Δ-3x-irradiated clone A6 | Illumina | This study |
| *PfRad51*Δ-3x-irradiated clone B4 | Illumina | This study |
| *PfRad51*Δ-3x-irradiated clone D1 | Illumina | This study |
| *PfRad51*Δ-3x-irradiated clone D7 | Illumina | This study |
| *PfRad51*Δ-3x-irradiated clone G4 | Illumina | This study |
| *PfRev1*Δ-3x-irradiated clone A5 | Illumina | This study |
| *PfRev1*Δ-3x-irradiated clone B7 | Illumina | This study |
| *PfRev1*Δ-3x-irradiated clone C4 | Illumina | This study |
| *PfRev1*Δ-3x-irradiated clone D4 | Illumina | This study |
| *PfPolζ*Δ-3x-irradiated clone A2 | Illumina | This study |
| *PfPolζ*Δ-3x-irradiated clone B5 | Illumina | This study |
| *PfPolζ*Δ-3x-irradiated clone B11 | Illumina | This study |
| *PfPolζ*Δ-3x-irradiated clone E5 | Illumina | This study |
| *PfPolζ*Δ-3x-irradiated clone F2 | Illumina | This study |
| *PfPolζ*Δ-3x-irradiated clone G8 | Illumina | This study |
| WT-*Upf1*Δ | Illumina | This study |
| *PfRad51*Δ-*Upf1*Δ | Illumina | This study |
| *PfRev1*Δ-*Upf1*Δ | Illumina | This study |
| *PfPolζ*Δ-*Upf1*Δ | Illumina | This study |
| WT-*Upf1*Δ-distant | Illumina | This study |
| *PfRad51*Δ-*Upf1*Δ-distant | Illumina | This study |
| *PfRev1*Δ-*Upf1*Δ-distant | Illumina | This study |
| *PfPolζ*Δ-*Upf1*Δ-distant | Illumina | This study |
| WT-*v2csa*Δ | PacBio | [3] |
| *PfRad51*Δ-*v2csa*Δ | Nanopore | This study |
| *PfRev1*Δ-*v2csa*Δ | Illumina | This study |
| *PfPolζ*Δ-*v2csa*Δ | Illumina | This study |

| | | |
|---|---|---|
| WT-v2csaΔ-distant | Nanopore | This study |
| *PfRad51*Δ-*v2csa*Δ-distant | Nanopore | This study |
| *PfRev1*Δ-*v2csa*Δ-distant | Nanopore | This study |
| *PfPolζ*Δ-*v2csa*Δ-distant | Nanopore | This study |

Supplementary Figure 1: *var* clusters in the 14 chromosomes of 3D7 *P. falciparum*

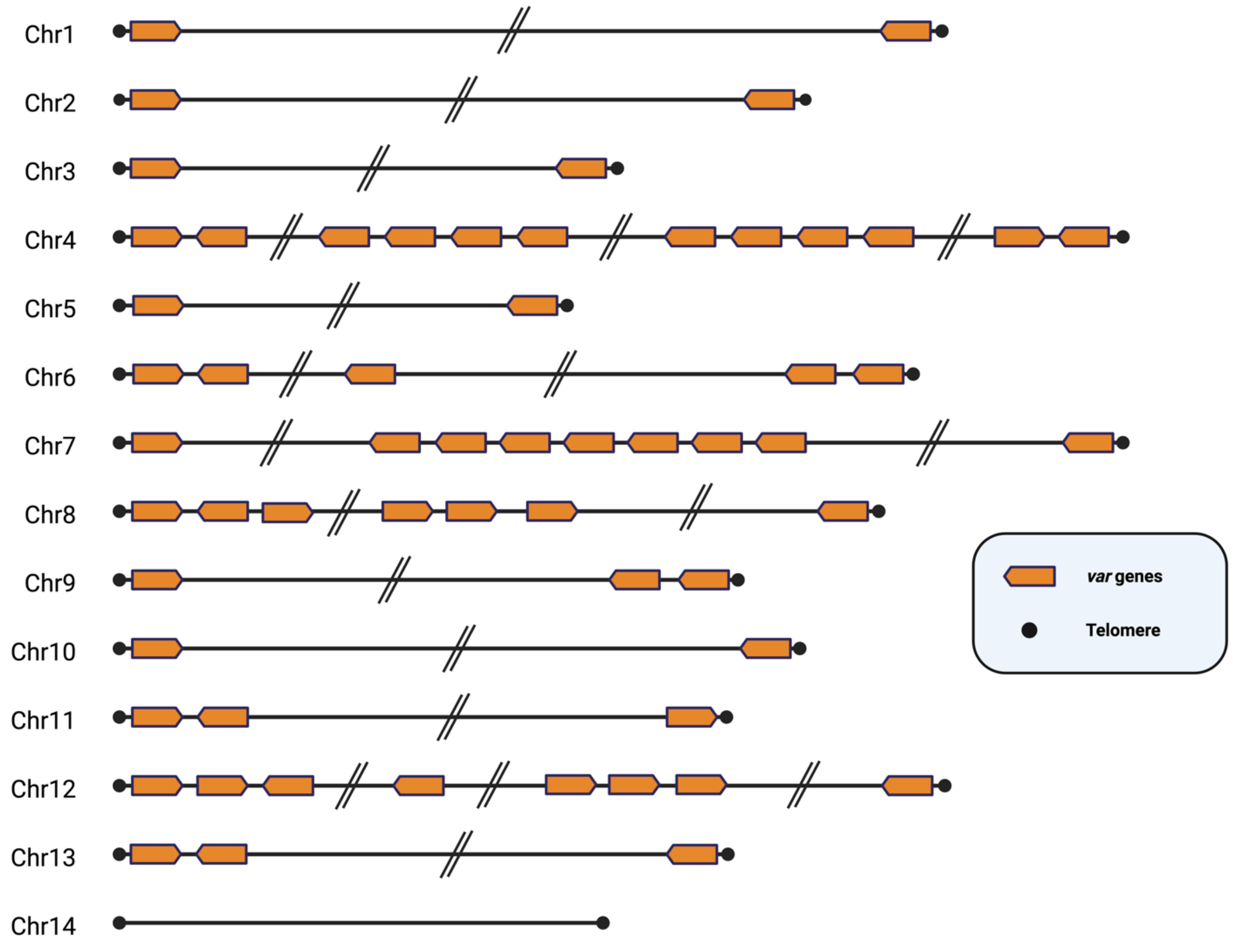

Supplementary Figure 2: Representative growth curves of non-irradiated knockout cell lines from this study. Cultures were grown from same initial parasitemia till they crashed. All the cell lines grew similar to the WT 3D7 reaching maximum parasitemia and crashing within 3-4 days.

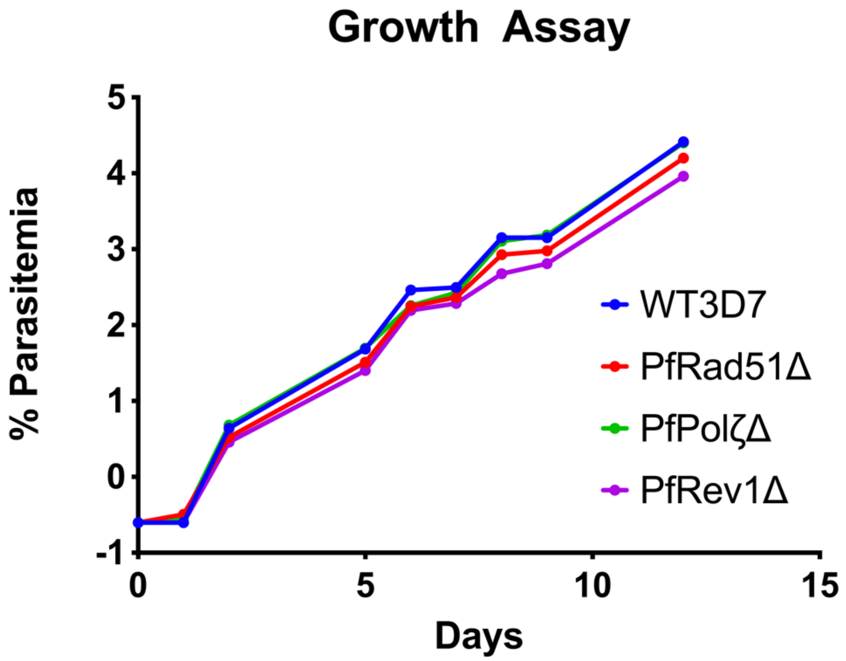

Supplementary Figure 3: Distribution of SNPs across genome: (A) Distribution of SNPs across the genome based on the location - Upstream and downstream of the coding regions, exons, introns and intergenic regions expressed as percentage. Statistical analysis is performed using One way ANOVA followed by Šídák's multiple comparison test. *($p$=0.01-0.05),**($p$=0.01-0.001),***($p$=0.001-0.0001).

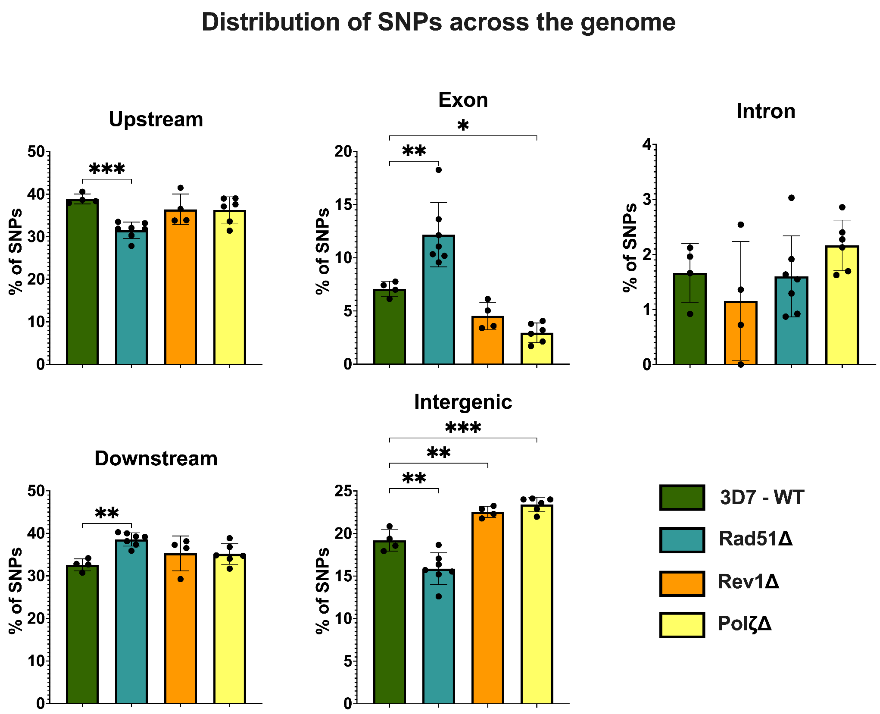

Supplementary Figures 4: IGV screenshots of Translocations in Irradiated WT 3D7 shown in Fig 5. The mismatches are highlighted with colors using the ‘show mismatch’ toggle in IGV. The red colored reads depict the forward reads while the blue colored reads are reverse reads. The black box indicates the SV observed in the chromosome and indicates the alignments that identified the breakpoint.

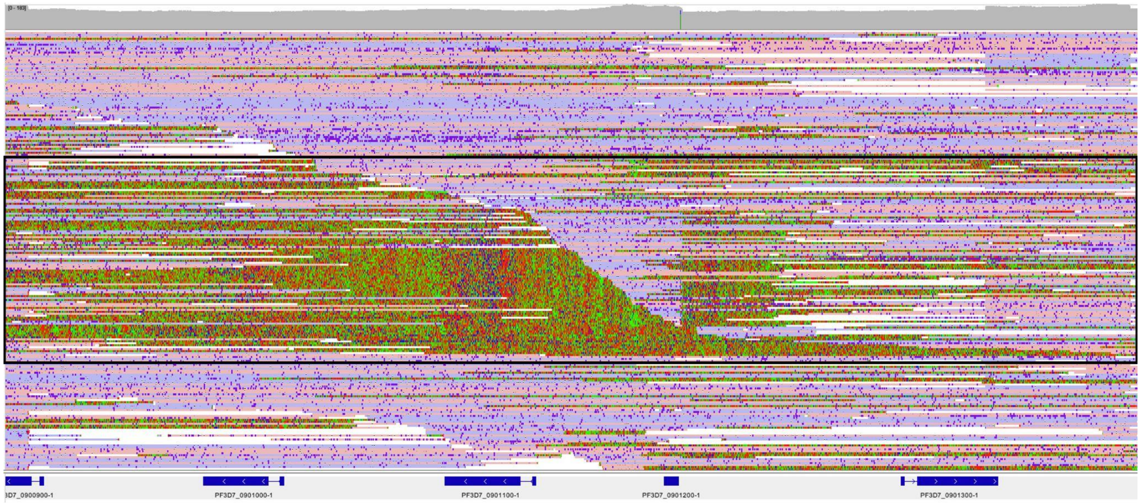

Chr14-Chr9 BND – Chr9 part; Clone F12

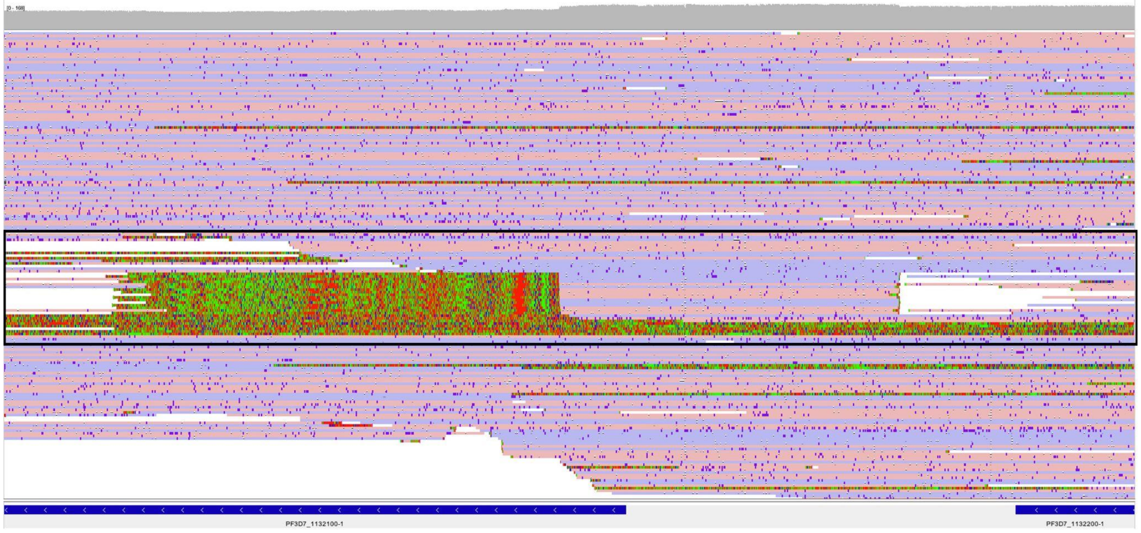

Chr11-Chr14 BND – Chr11 part ; Clone F12

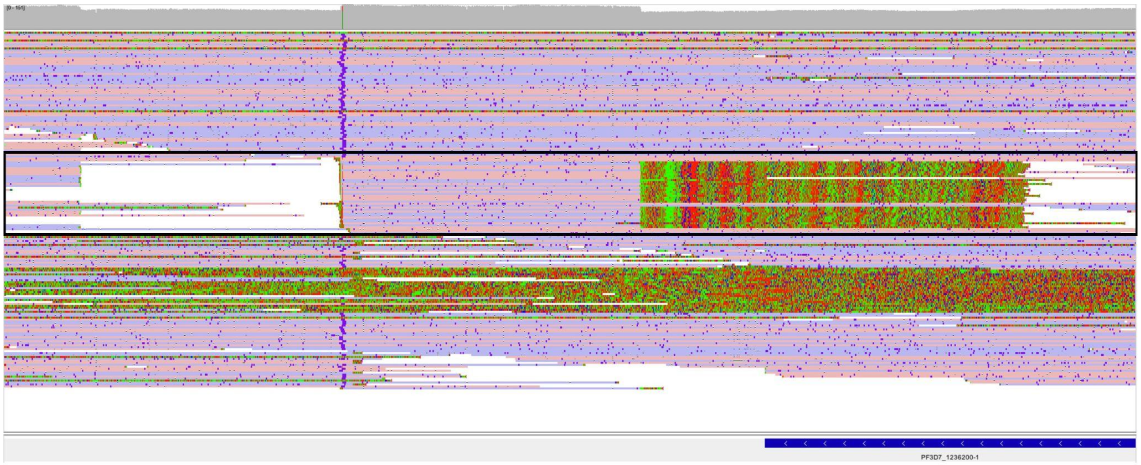


Chr12-Chr13 BND – Chr12 part ; Clone F12

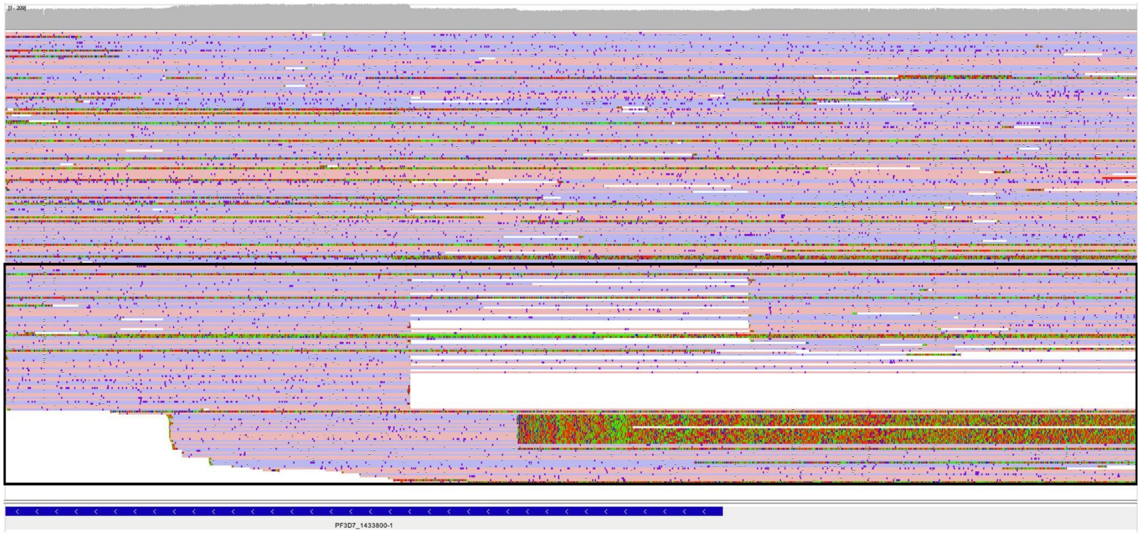


Chr14-Chr5 BND – Chr14 part ; Clone F12

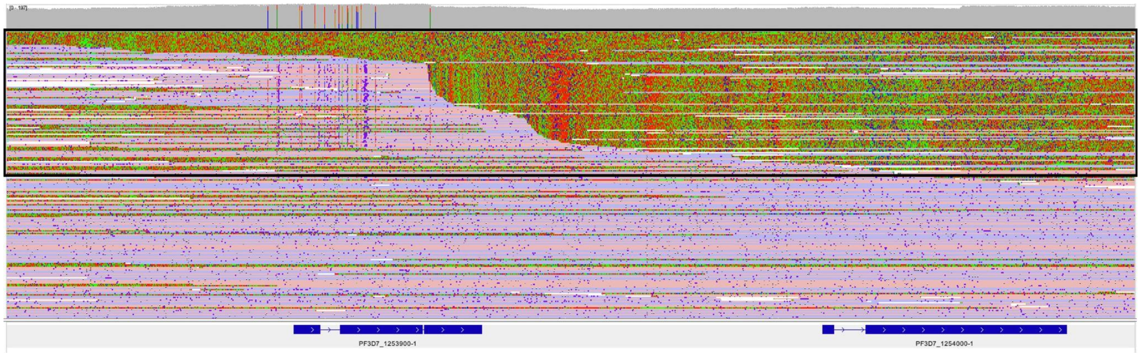


Chr12-Chr9 BND – Chr12 part ; Clone F12

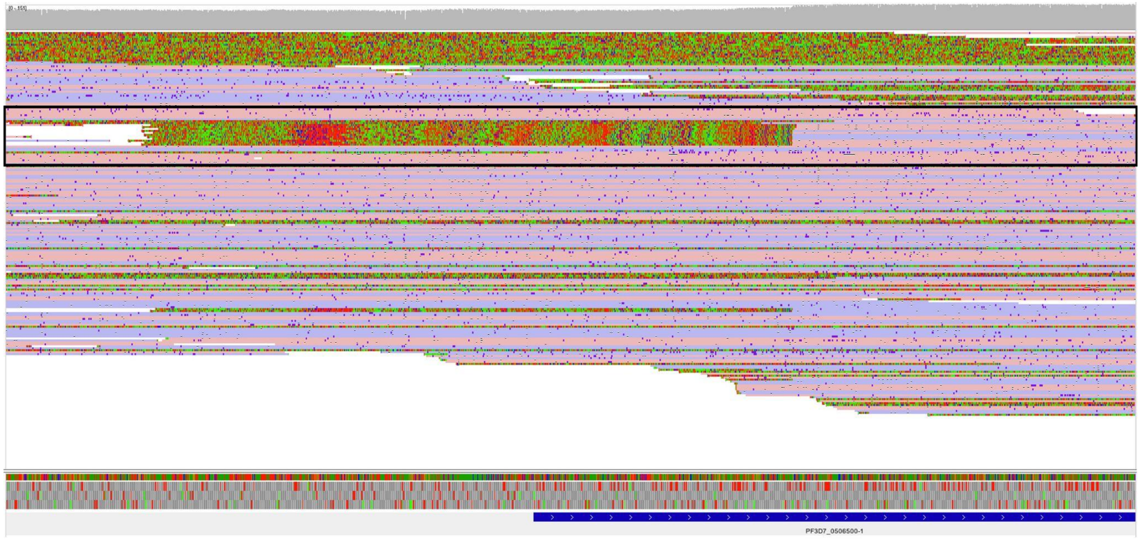


Chr5-Chr8 BND – Chr5 part ; Clone F12

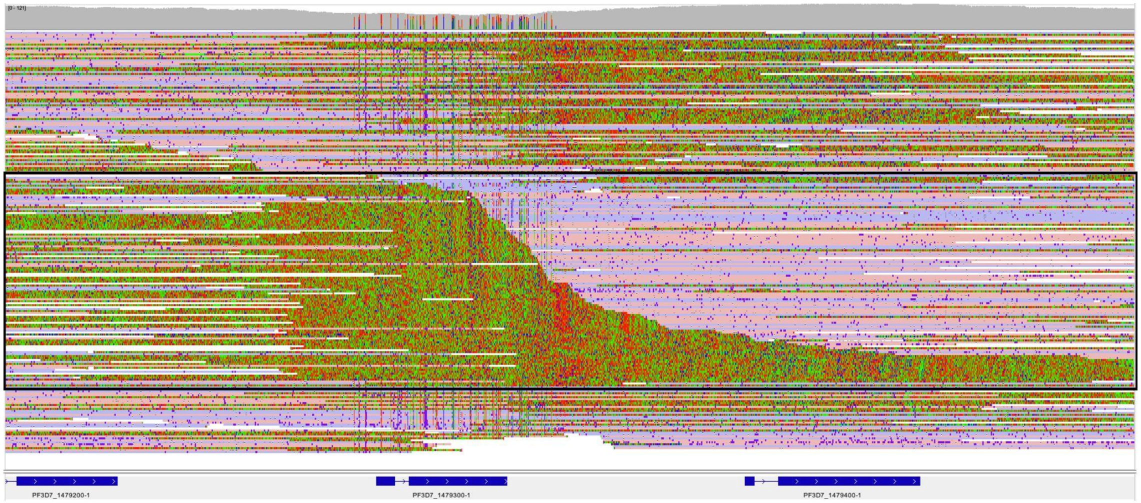


Chr14-Chr9 BND – Chr14 part ; Clone F12

Supplementary Figure 5: IGV screenshot of the read pileup of *PfRad51*Δ-D9 clone at the EMP1-trafficking protein locus showing a large deletion in the repetitive sequence blocks. The red colored reads depict the forward reads while the blue colored reads are reverse reads.

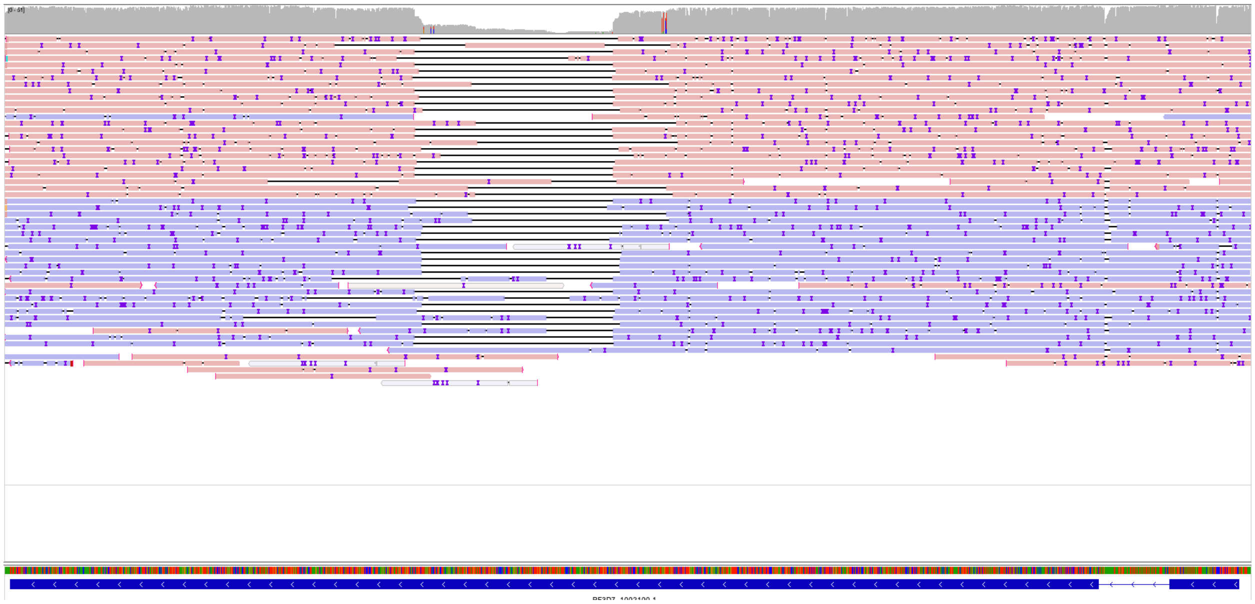

Supplementary Figure 6: IGV screenshots of the read pileup of distant knockout constructs. The mismatches are highlighted with colors using the 'show mismatch' toggle in IGV. The red colored reads depict the forward reads while the blue colored reads are reverse reads.

PfRad51Δ-var2csaΔ-distant

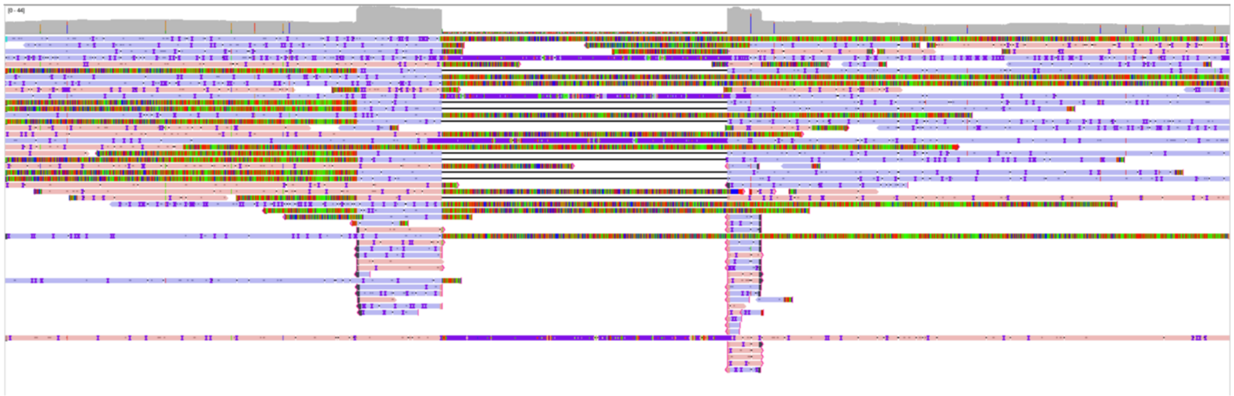

PfRev1Δ-var2csaΔ-distant

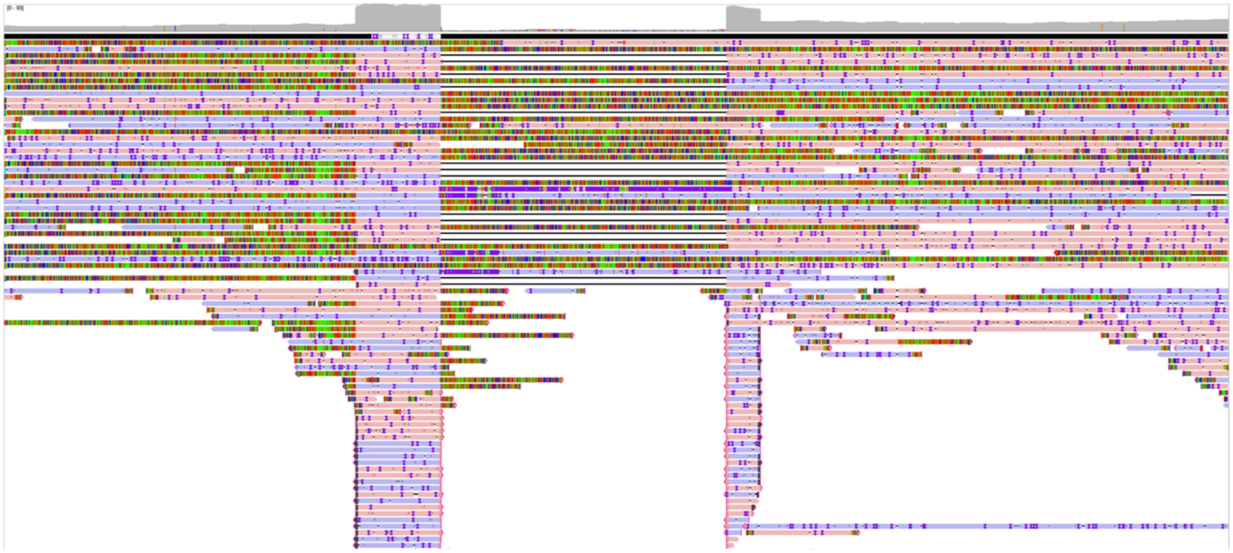

PfPolζΔ-var2csaΔ-distant

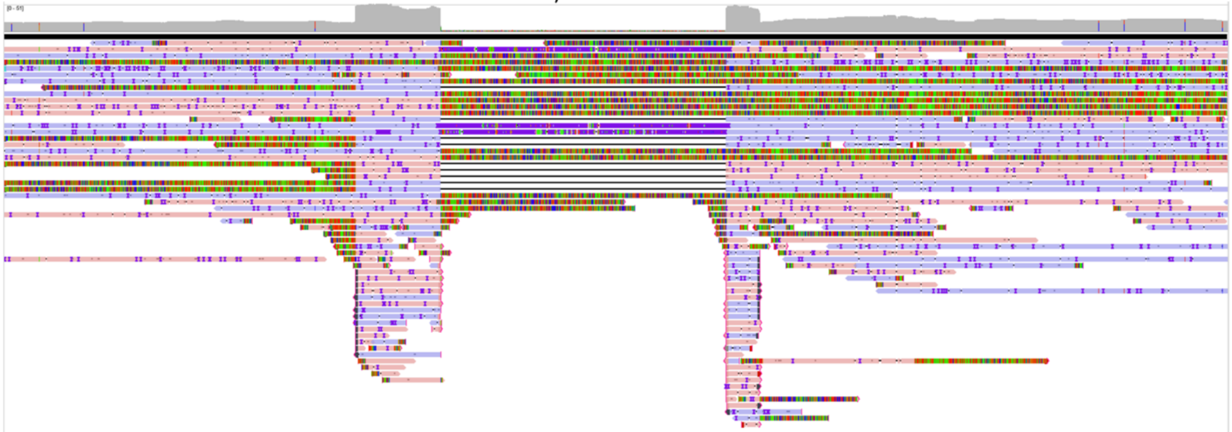
